# StrucNS reveals interaction-weighted network topology as the driving predictor of absolute stability of natural and de novo proteins

**DOI:** 10.64898/2026.05.25.727723

**Authors:** Archita Mullick, Prodromos Daoutidis, Benjamin J. Hackel

## Abstract

**Motivation:** Folded protein function requires stability, yet mapping structure and sequence to a fitness landscape remains difficult. The protein fold is the physical realization of complex, spatially-sensitive physicochemical interactions among residues; quantitatively elucidating how these subtle relationships dictate thermodynamic stability remains challenging. We present StrucNS, a mathematical framework that identifies principles governing protein fitness by employing network science to learn physicochemical relationships between residues and their stability contributions directly from the protein fold. Representing the fold as a network topology, we utilize an inverse approach: while the fold is traditionally viewed as the phenotypic consequence of the underlying chemical forces, we use the topology to decode the very physicochemical dependencies that govern protein stability. Unlike protein language models reliant on high-dimensional evolutionary embeddings, StrucNS extracts these signals directly from the interaction-weighted network topology. Independence from evolutionary history uniquely suits StrucNS for de novo design prediction.

**Results:** Despite reduced dimensionality and training depth, StrucNS outperforms ESM-2 and ProteinMPNN on predicting mutational stability. StrucNS outperforms supervised UniRep in predicting absolute stability of de novo designs. Feature analysis reveals network topology as the key driver of predictive power, contributing 59% of model importance. SHAP analysis reveals two highly influential features as high degree and low modularity of polar/hydrophobic mixed subnetworks, which highlights the importance of connectivity between the hydrophobic core and protein surface to drive stability contrary to the conventional focus on the hydrophobic core. Revelation of predictive topological features underscores the utility of an interpretable model.

**Availability:** Source codes are available: https://github.com/Hackel-Group-CEMS/StrucNS.

## Introduction

Proteins represent the ultimate frontier of programmable matter, serving as the primary functional scaffolds that orchestrate the complexity of biological life [1, 2]. From the precision of intracellular signal transduction to the robust mechanics of systemic immunity, the utility of these macromolecules is derived from their ability to adopt unique three-dimensional architectures [3]. Beyond their endogenous roles, the capacity to rewrite protein sequences to provide improved or unique function is the cornerstone of modern biotechnology, enabling the synthesis of targeted therapeutics, high-sensitivity diagnostics, and industrial biocatalysts [4, 5]. At the heart of these advancements lies the physical requirement, for folded proteins, that a linear polypeptide sequence must not only fold into a specific geometry but also possess the thermodynamic stability necessary to resist the entropic drive toward unfolding [6, 7] all while enabling functionality (e.g. binding or catalysis). Understanding the determinants of this stability empowers protein engineering and is a prerequisite for interpreting the functional consequences of natural genetic variation [8].

Despite decades of research, the protein fitness landscape remains a *terra incognita* of immense dimensionality. The spatial geometry of a fold is the physical manifestation of many-body physicochemical interactions—a complex interplay of hydrophobic packing, hydrogen-bonding networks, and configurational entropy [9, 10]. However, the precise mapping between 3D coordinates and the underlying energy landscape remains an enduring enigma of molecular biophysics. Traditional experimental paradigms, such as directed evolution, attempt to traverse this landscape via stochastic sampling [11]. While powerful, these methods are fundamentally constrained by the “curse of dimensionality”, as they can only explore a small fraction of the potential 20^*N*^ sequence space—a space larger than the number of atoms in the observable universe [12]. Conversely, rational design seeks to apply structural priors to introduce targeted mutations, yet our fragmentary understanding of the non-linear dependencies and the subtle topological constraints and geometric tensions within the native state often results in unreliable heuristics, limiting the success of classical force-field modeling [13].

As an alternative approach, machine learning has emerged as a transformative framework for inferring the protein energy landscape. Modern deep-learning architectures, including protein language models and graph neural networks, have demonstrated an unprecedented ability to learn the statistical regularities inherent in vast biological datasets, acting as high-throughput digital surrogates for select laboratory assays [14, 15]. However, as these models evolve in complexity, a fundamental interpretability crisis has emerged: a trade-off between predictive accuracy and physical transparency [16, 17]. While these black-box architectures achieve high empirical accuracy, they offer little insight into the structural logic governing their predictions. For instance, in the design of *de novo* proteins, which lack the evolutionary history that modern models often rely upon, an opaque model may correctly flag a candidate as unstable without identifying the specific topological defect responsible [18]. This lack of mechanistic feedback creates a bottleneck in the design cycle; without knowing *why* a fold fails, the engineer cannot inform the next round of sampling, effectively stalling the discovery pipeline. Consequently, there is a pressing need for a framework that transcends pattern matching to recover the fundamental structural rules that drive protein stability.

A dichotomy currently exists in protein modeling between zero-shot inference and high-capacity supervised learning. Zero-shot models have fundamentally redefined our ability to navigate sequence space by treating proteins as a structured language, learning the “evolutionary grammar” from vast genomic repositories [19, 20]. By distilling statistical patterns of conservation and co-variation, sequence-centric models — most notably protein language models (PLMs) such as ESM-2 — capture a global summary of biological plausibility, enabling the prediction of mutational effects across diverse protein families without stability-specific training [21, 14]. Within this paradigm, the emergence of ProteinMPNN has established a new benchmark for structure-based zero-shot design [22]. By utilizing a deep message-passing neural network to learn the conditional probability of an amino acid sequence given a fixed three-dimensional backbone, ProteinMPNN effectively captures the local packing requirements and steric constraints of the native state [15]. However, despite its generative prowess, ProteinMPNN is primarily optimized for the inverse folding problem; it identifies sequences that are *compatible* with a fold but does not explicitly quantify the energetic magnitude of thermodynamic stability or the biophysical drivers of destabilization.

To achieve the high-resolution precision required for biochemical applications, researchers have increasingly turned to one-shot and supervised learning frameworks. In these paradigms, the high-dimensional latent representations of PLMs, such as the 1280-dimensional embeddings of ESM-2 or the 1900-dimensional hidden states of UniRep, are recycled as informative priors for specialized downstream tasks [23]. For instance, UniRep embeddings have been successfully utilized to guide the optimization of green fluorescent proteins and industrial enzymes with minimal experimental data [20, 24]. Building on this logic, ThermoMPNN and the SPURS (Stability Prediction Using a Rewired Strategy) framework represent the state-of-the-art in integrating structural and evolutionary signals [25]. ThermoMPNN combines the structural encoding of ProteinMPNN with a supervised regressor trained on Megascale dataset [26] of 272,712 variants to map structural features directly to changes in the Gibbs free energy of unfolding (ΔΔ*G*) [25]. SPURS further pushes this synergy by “rewiring” the structural features from ProteinMPNN into the attention layers of ESM-2, effectively conditioning evolutionary signals on 3D geometric constraints to predict absolute stability.

Despite the power of these synergistic frameworks, a critical generalization ceiling and interpretability gap remain. While PLMs like ESM-2 and UniRep are powerful, their heavy reliance on evolutionary depth makes them prone to failure when applied to *de novo* proteins or orphan sequences that lack a rich history of natural selection [18, 27]. Recent studies have highlighted the limitations of sequence-only models in navigating the “fitness deserts” of synthetic proteins, where the rules of biological plausibility are not yet established by evolution [28]. Furthermore, the reliance on dense latent embeddings creates a strategic bottleneck in the design cycle. Because these architectures operate within an opaque mathematical manifold, they lack the physical transparency required for informed engineering [29]. When a black-box model rejects a candidate library, it offers no actionable mechanistic feedback. Post-hoc explanation methods like SHAP (SHapley Additive exPlanations) have been shown to be fragile in these settings, with embedding-based attributions often failing to correspond to discrete biophysical entities such as salt bridges or hydrogen-bond networks [30, 31]. This lack of mechanistic grounding creates a trust gap in rational engineering. Consequently, there is a necessity for a framework that does not merely match patterns in a latent space but recovers the fundamental, interpretable structural rules that drive protein stability.

We propose that this bridge can be built through a network topology-based abstraction: conceptualizing the protein not as merely a fold geometry, but as an interaction-weighted network topology. This shift addresses a central challenge in structural biology: the realization that protein fold is the physical manifestation of hidden, multi-scale interactions [32, 33]. By applying this perspective at the Ångström scale, we can treat thermodynamic stability as an emergent property of the network topology, allowing us to test a fundamental hypothesis: that defined topological invariants can serve as sufficient surrogates for the intricate forces governing the fitness landscape [34].

To investigate this, we present StrucNS, a framework designed to decode the physicochemical logic of stability of a fold through a topological lens. Unlike existing models that rely on high-dimensional evolutionary embeddings [35, 36], StrucNS operates as a “no-embedding” framework, deriving its predictive power directly from the explicit connectivity of a residue interaction network (RIN) [37, 38, 39]. The first pillar of our approach focuses on the hierarchical organization of the fold. We partition the protein into discrete topological subnetworks based on chemical environments, such as the hydrophobic core and polar surface, to analyze the balance between connectivity within a community and the interactions across these boundaries. This allows us to determine whether the predictive signal for stability can be recovered directly from the topological organization of these communities, offering a pathway to evaluate *de novo* protein variants where evolutionary history, the primary signal for current machine learning models, is absent or poorly defined [40, 41].

To validate the generalizability of this topological logic, we benchmarked StrucNS against diverse mutational datasets, spanning both natural folds and synthetic designs. StrucNS outperforms PLMs in predicting mutational stability, reveals the value of a network science perspective on modeling protein structure/stability relationships, and highlights an interaction-weighted network topology as a key determinant of protein stability prediction. StrucNS outperforms ESM and MPNN models in predicting mutational stability, especially in de novo folds. StrucNS performs similarly to supervised Unirep and is competitive with supervised ESM-2 despite reduced model dimensionality and training depth. Crucially, despite being trained exclusively on miniproteins, StrucNS demonstrates generalizability, achieving performance on large globular proteins over 10 times the size of the folds in the training set.

StrucNS offers a transparent, biophysically grounded lens through which to understand and engineer the absolute stability of both natural and de novo proteins. Our analysis reveals that network topology is the primary driver of protein stability, as evidenced by 59% feature importance and 53% recall reduction upon topology feature ablation. Intriguingly, our findings reveal that the hybrid polar/hydrophobic subnetworks are more influential than the hydrophobic core in determining protein stability, with high connectivity and low modularity as stabilizing features. By unifying these topological insights with high-throughput stability prediction, StrucNS demonstrates that the precise architecture of energy-weighted residue interactions is a fundamental determinant of protein fitness, providing a powerful and interpretable framework for the computational design of robust, highly stable proteins.

## Methods

### StrucNS architecture

We hypothesize that thermodynamic stability, in both natural and de novo proteins, is an emergent property of hierarchical interaction topology. To test this, we developed StrucNS, a multi-scale framework that decodes high-dimensional spatial data into interpretable topological signals. For clarity, we define the following nomenclature: the protein fold refers to the explicit three-dimensional spatial coordinates of the residues; network topology represents the protein as an abstract graph, defined by its nodes (residues), edges (interactions), and spatial positioning.

StrucNS (Fig. 1) operates through network science operations: (i) construction of an energy-weighted RIN from in-silico structures [42, 3]; (ii) identification of mesoscopic partitions of dense interactions through community detection; (iii) identification of macroscopic subnetworks through functional semantic labeling; (iv) mechanical equilibrium of an interaction-weighted network; (v) calculation of network features; and (vi) model training.

**Figure 1.**
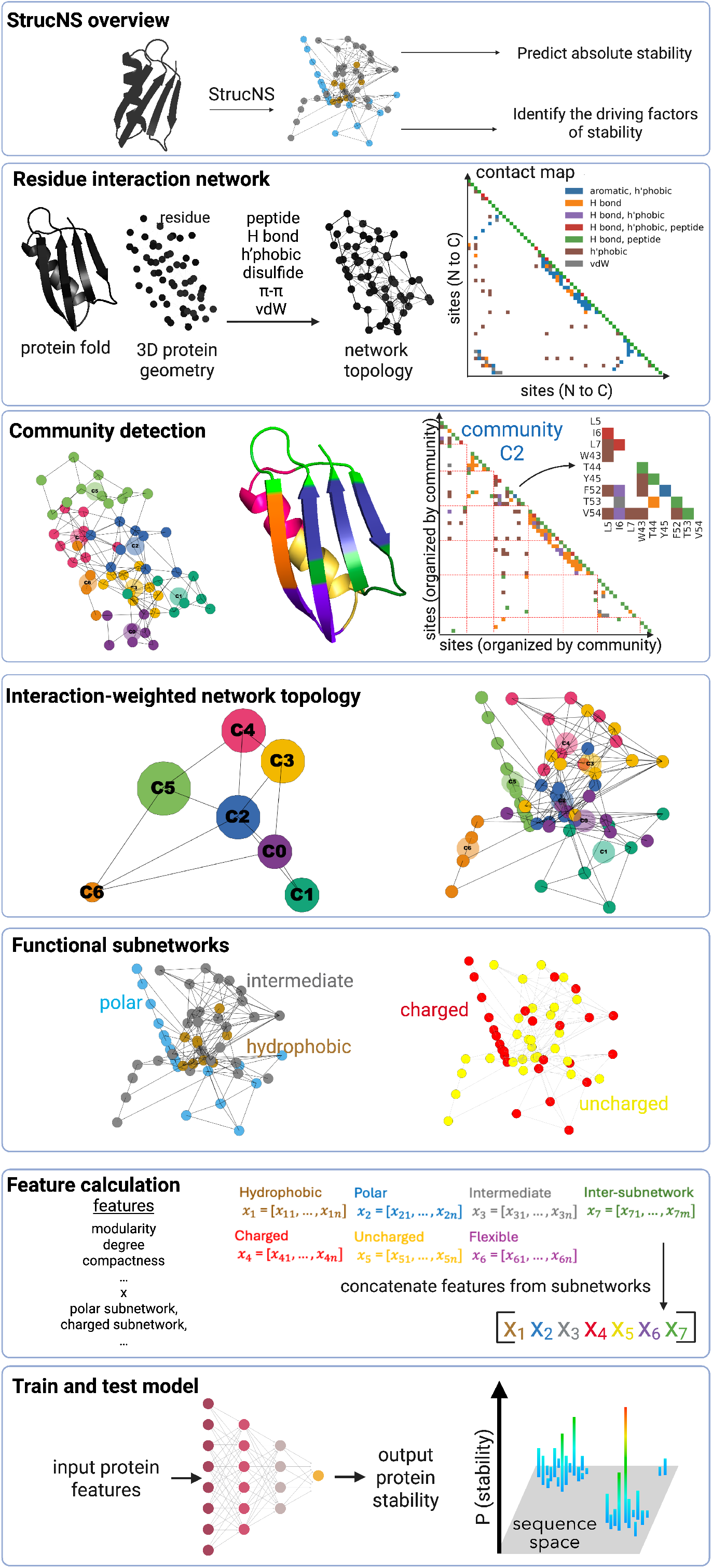
Overview of StrucNS model. StrucNS uses protein folds to build residue interaction networks and community-driven network topology, allowing it to predict absolute thermodynamic stability while feature analysis enables determination of the driving factors of protein stability. A *ββαββ* protein (PDB: 1PGA) is used to exemplify the StrucNS approach. *Residue interaction network (RIN)*. The three-dimensional protein fold is converted into a discrete RIN in which each amino acid centroid is a node and six bond types are identified as edges based on spatial and physicochemical criteria. These interactions are shown as a contact map, with white being not connected and color being a type of interaction. *Community detection*. Communities are then detected as clusters of nodes based on modularity maximization. As shown in the community contact map, these well-connected clusters appear as dense blocks along the diagonal, while the rest of the matrix remains sparse, highlighting the modular nature of the identified communities. Community 2 (C2) is enlarged as an example to show its members (residues 5, 6, 7, 43, 44, 45, 52, 53, 54). *Interaction-weighted network topology projection*. Node positions are then adjusted by performing a spring balance with attractive edges and repulsive non-edges to generate an interaction-weighted network topology projection. *Functional subnetworks*. Communities are then labeled and grouped into functional subnetworks on the basis of physicochemical composition (polar, intermediate, hydrophobic; charged or uncharged). *Features estimation*. The labeled topology is then evaluated for a series of network science features within and across subnetworks. *Train and test model*. This collection of features is then used as inputs to train a mathematical model to predict protein stability as the output across a broad swath of protein structures and sequences. The resultant model enables a map of the sequence-stability landscape.

### Residue interaction network (RIN)

Amino acid sequences were converted to FASTA format and processed using OmegaFold (v1.1.0). Structure predictions were performed using the 60M-parameter model with default weights and four cycles. Computations were executed on NVIDIA A100 GPUs with a 15-minute runtime limit per sequence. We modeled the protein tertiary structure as an energy-weighted RIN using the `Biopython` (v1.81) PDB parser and `NetworkX` (v3.1). Nodes in the network represent residue centroids, serving as the effective centers of interaction mass for each side chain (Fig. 1). Edges represent specific physicochemical interactions: covalent peptide bonds, hydrogen bonds, hydrophobic contacts, disulfide bridges, aromatic stacking, and van der Waals forces. We established these edges using stringent atomic distance thresholds, such as a 3.5 Å limit for hydrogen bonds between donor and acceptor atoms of neighboring residues. Detailed criteria are provided in SI Appendix, S1.1. To reflect disparate thermodynamic contributions, we weighted edges using established bond dissociation energies and experimental stability scales (SI Appendix, S1.2). Fig. 1 *RIN* illustrates the protein’s tertiary structure and network topology, including residue coordinates and edge weights. The resulting connectivity map highlights the network’s sparseness and the diverse interaction types between residues.

### Community detection via weighted modularity optimization

The RIN provides a high-resolution map of atomic interactions, yet the resulting topology is inherently high-dimensional, as seen in the contact map (Fig. 1, *RIN*). To mitigate structural noise common in structure prediction models like OmegaFold [42], we transitioned from a single-node representation to a coarse-grained, mesoscopic framework. We applied a network partitioning strategy based on modularity maximization to identify discrete community structures (Fig. 1, *Community detection*). In this context, we hypothesized that because RIN edges and weights reflect physical chemical interactions, modularity optimization would isolate “interaction-dense” communities. These clusters represent residue groups where the internal density of biochemical bonds significantly exceeds the connectivity expected from a random null model. This null model serves as a critical statistical baseline, representing stochastic connectivity while preserving the degree of each residue (SI Appendix, S1.3). Using the Louvain algorithm [43] on the energy-weighted RIN, we decomposed *N* protein residues into *K* discrete structural modules (*K < N*). Notably, these communities frequently incorporate residues distant in primary sequence. For instance, Community 2 (C2) spans disparate secondary structures, including multiple *β*-sheets (Fig. 1 *Community detection*). This partitioning remains entirely network-driven and agnostic to specific residue identities. These communities, marked by triangles along the contact map diagonal, exhibit higher internal connectivity than interactions with the rest of the network.

### Interaction-weighted network topology via spring balance

Protein tertiary structure arises from the interplay between short-range packing forces within a secondary structure and long-range interactions across secondary structures. To encode this information, we transformed the protein fold into an interaction-weighted network topology by treating the network topology as a collection of springs. Connected nodes experienced an attractive force proportional to the edge weight, while unconnected nodes experienced a repulsive force proportional to the inverse of 1*/*4th of their distance. We applied a force balance (SI Appendix, S1.4) on the nodes using the Fruchterman-Reingold algorithm [44] to reach a topological equilibrium (Fig. 1, *Interaction-weighted network topology projection*). This new projection represents a state of topological equilibrium that embeds interactions into the projection space. First, we applied a spring-balance to this global topology to rearrange nodes and communities according to equilibrium. Next, we applied spring-balance to the intra-community nodes until they reached equilibrium. We performed this in two steps based on the hypothesis of resolving local and global level forces. Models in literature, such as AlphaFold3 [45], have achieved success by resolving local and global scale interactions, which informed our choice of this hypothesis.

### Macroscopic subnetworks through functional semantic labelling

Modularity optimization partitions the protein into structurally coherent units based on interaction density. However, these emergent communities are initially defined by arbitrary numerical indices that lack biochemical context. To bridge mathematical interaction-weighted network topology with the thermodynamic principles governing protein stability, we implemented multi-functional semantic labeling (SI Appendix, S1.6) to identify functional subnetworks (Fig. 1, *Functional subnetworks*). While community detection identifies clusters of densely interconnected residues, semantic labeling defines the “biochemical personality” of these modules. This process characterizes how each functional subnetwork contributes to the global stability of the protein. We hypothesize that communities are biophysically pleiotropic; a single community may exert multiple, simultaneous functions on the fold. For instance, a rigid hydrophobic community may concurrently harbor charged residues at its solvent-exposed surface, fulfilling dual roles. To capture this complexity, we labeled each community using a composite hydrophobicity metric (derived from normalized Kyte-Doolittle [46] and Grantham scales [47]), charge metrics, and flexibility thresholds (SI Appendix, S1.6). We grouped communities with shared labels into three primary subnetworks: hydrophobic, polar, and intermediate (Fig. 1, *Functional subnetworks*). For example, Community 4 (C4) belongs to both the intermediate (grey) and charged (red) subnetworks, and Community 3 (C3) resides within both the hydrophobic (brown) and uncharged (yellow) subnetworks.

### Functional subnetwork-based and inter-subnetwork feature extraction

By this stage, we have established an interaction-weighted network topology partitioned into overlapping functional subnetworks. We now translate this architecture into a feature space, defined as a vector of parameters describing network science features of individual functional subnetworks (e.g. modularity of each polar, hydrophobic, etc.) and inter-subnetworks (e.g. boundary flux between polar and hydrophobic) (full feature list and descriptions in SI Appendix, S1.7). These features derive three types of information: connectivity-based, protein fold geometry-based, and interaction-weighted network topology-based. Connectivity-based features depend solely on nodes and edges, including modularity, the density of internal connections relative to a random null model, edge density, the ratio of actual edges to all possible connections, and degree, the number of connections per residue. Fold geometry-based features depend on the distribution of residues in 3D space. Interaction-weighted network topology-based features inform the interaction-weighted geometry of subnetworks, such as the spatial clustering coefficient of each subnetwork, which quantifies the local node density within the interaction-weighted space. Once we estimate these features from each of the six subnetworks and the inter-subnetwork features, we concatenate them into a single vector to form the feature space of the protein (Fig. 1, *Feature calculation*).

### Training and Test Sets

To construct a robust predictive framework, we utilized a dataset with absolute protein stability measured, via cDNA display proteolysis, for 844425 natural and de novo proteins [26]. The Megascale dataset was downloaded from Zenodo (https://zenodo.org/records/7992926). We filtered the primary dataset (`Tsuboyama2023_Dataset1_20230416.csv`) to retain datapoints which had confidence interval lower than ∼1.6 kcal/mol to ensure that the target labels, representing the experimental change in Gibbs free energy, originated from high-quality thermodynamic measurements. To rigorously evaluate whether our model learns universal physicochemical principles or simply memorizes specific backbone geometries, the data were partitioned into three mutually exclusive test sets based on protein family identity and mutational coverage (Fig. 2A). The data were divided such that 80% of the available variants from 80% of parental folds were used for model training. The remaining 20% of variants from those same parental folds were set aside as Test Set 1 (TS1). This internal validation set measures the model’s ability to interpolate within known structural architectures by predicting the stability of unseen mutational variants on scaffolds that were already presented during the optimization phase. To evaluate true structural generalization, we established two additional external holdout sets, Test Set 2 (TS2) and Test Set 3 (TS3), which contain protein families that were entirely excluded from the training process. These sets represent unseen folds that the model has never encountered during its optimization. TS2 comprises de novo designed proteins—synthetic structures engineered by researchers to adopt specific folds, while TS3 consists of naturally occurring parental folds. Success on TS2 and TS3 would demonstrate a capacity to generalize structural logic across the structural universe and prove that the model can interpret the stability of entirely novel architectures rather than relying on the specific geometric features of the training examples.

**Figure 2.**
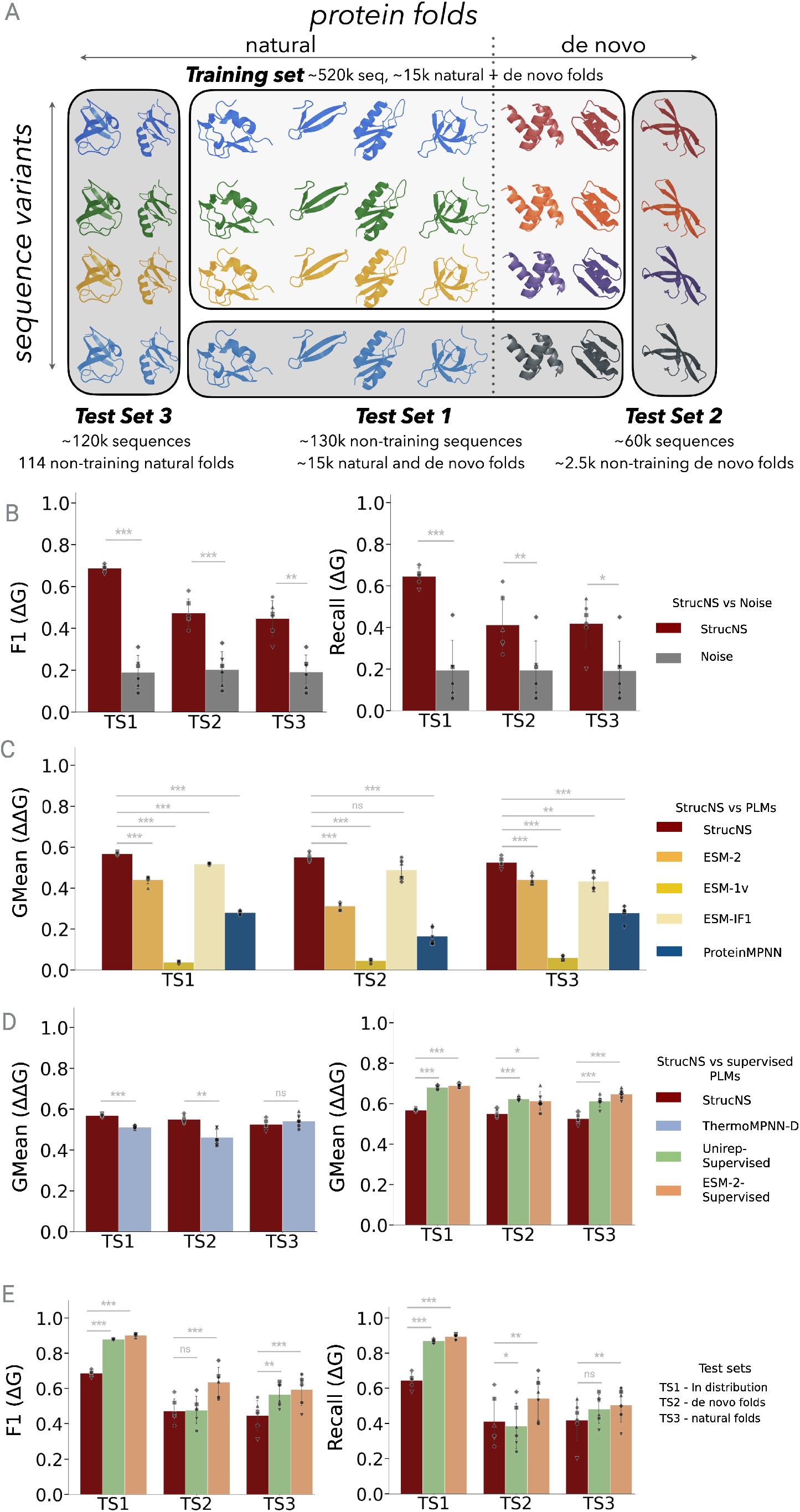
The predictive performance of StrucNS across broad test sets (A) was evaluated in comparison to (B) statistical noise, (C) evolutionary models, and (D, E) high-capacity embeddings in supervised protein language models. StrucNS was trained on 80% of sequences within 80% of protein folds within the 8x10^5^ sequence-stability measurements in Megascale data set [26]. (A) Data are presented for evaluation across three test sets: TS1: sequences held out of training from parental folds included in model training; TS2: de novo designed proteins with parental folds excluded from training; TS3: natural protein folds excluded from training. Data are shown for each of six independent train/test splits (symbols) along with the mean and standard deviation. Statistical significance relative to StrucNS – as computed via paired t-test – is depicted with * (0.01 *<* p *≤* 0.05), ** (0.001 *<* p *≤* 0.01), and *** (p *≤* 0.001). (B) The recall (left) and F1 statistic (right) for predicting the stability classification (ΔG *>* 3 kcal/mol) of variants is shown for StrucNS and a noised model in which feature values were randomly permuted across all variants, thereby preserving the feature distribution but ablating connection to the proper stability output. (C) The geometric mean of classwise sensitivity (GMean) for predicting the proper change (increase or decrease) in the probability of stability upon mutation was computed using StrucNS or protein language models ESM-2 [21], ESM-1v [14], ESM-IF1 [27], or ProteinMPNN [22]. (D) The GMean for predicting mutational stabilization was computed using StrucNS or ThermoMPNN [25] (left), which was trained on the same Megascale data set, and supervised implementations of UniRep or ESM-2 (right). (E) Recall and F1 for predicting the stability classification of variants was computed for StrucNS and the supervised protein language models.

### Performance metrics

The predictive objective was formulated as a binary classification task to distinguish robustly stable protein variants from unstable or marginally stable backgrounds. While thermodynamic stability is conventionally bounded at a free energy of unfolding, Δ*G >* 0, we imposed a stricter classification threshold of Δ*G >* 3 kcal/mol. This rigorous cutoff was selected to isolate variants with high structural resilience, effectively filtering out neutral drift and ensuring that “stable” classifications correspond to designs with significant thermodynamic utility. In protein engineering workflows, downstream experimental characterization represents the primary bottleneck; therefore, we assessed precision to minimize the resource burden of validating false positives. Recall was simultaneously monitored to ensure the computational filter maintained sufficient coverage of the fitness landscape, preventing the exclusion of diverse, viable candidates. We use the harmonic mean of these two metrics, F1, as a key merged metric. To further evaluate performance in the context of mutational stability, we employed the geometric mean (GMean) of sensitivity and specificity, a metric that provides a balanced assessment of classification accuracy across imbalanced datasets. The StrucNS model generates a likelihood of stability (Δ*G >* 3 kcal/mol), from which we derived a log-likelihood ratio (LLR) to compare each variant against the wild-type (WT) sequence. Variants with an LLR *>* 0.05 were classified as stabilizing mutations, while those with LLR *<* −0.05 were categorized as destabilizing. By utilizing GMean as a primary benchmark for mutational stability, we ensured a robust comparison of our approach against existing computational models.

### Machine learning model

To predict stability, we developed a binary classification model where samples were labeled as stable for Δ*G >* 3 kcal/mol and unstable for Δ*G* ≤ 3 kcal/mol. The architecture consists of a fully-connected feed-forward neural network implemented in TensorFlow, featuring a dynamic configuration of 1 to 5 hidden layers with widths ranging from 16 to 512 units. To ensure the robustness and generalization of our results, we generated six independent training/test cases, each utilizing a unique random seed to partition the full dataset into training, TS1, TS2, and TS3. This multi-split approach ensures that model performance is not an artifact of a specific data distribution. For each of these six cases, a dedicated model was developed through a rigorous hyperparameter optimization pipeline. Hyperparameter tuning for each case was conducted using the Optuna framework via 5-fold cross-validation. The predictive framework was implemented using `TensorFlow` (v2.12) and `Keras` . To optimize the neural network architecture, we utilized the `Optuna` (v3.2) framework, executing 300 trials managed via a `SQLAlchemy` persistent SQLite backend (`optuna study.db`). The objective function targeted the maximization of the average peak *F*_1_-score across all five folds (SI Appendix, S2, Fig. S2). To maintain the integrity of the validation process, a StandardScaler was fitted strictly on the training folds and applied to the validation fold to prevent data leakage. We also implemented fold-wise splitting for 5-fold cross validation, ensuring all sequences belonging to the same fold were grouped within the same fold to avoid overestimating accuracy through sequence similarity. The optimization objective maximized the *F*_1_ score using the `optuna.trial` interface to dynamically sample architectural parameters: number of hidden layers (1-5), layer units (`suggest_categorical`), `Dropout` rates (0.1-0.5), and `learning_rate` (logarithmic scale 10^*−*4^ to 10^*−*1^). Data preprocessing was handled through `scikit-learn`, employing `StandardScaler` to normalize feature distributions. To prevent data leakage, scaling parameters were computed strictly on training folds and applied to validation/test sets via the `fit transform` and `transform` methods. Cross-validation was managed through `sklearn.model_selection.KFold` with 5 splits. Model performance was monitored using a custom `keras.callbacks.Callback` to track *F*_1_ scores at each epoch end, while `EarlyStopping` was implemented to monitor `val_loss` with a patience of 8-10 epochs. The final model weights were selected based on the peak validation *F*_1_ score while monitoring validation loss. Statistical evaluation was performed using `sklearn.metrics` to compute precision, recall, and confusion matrices. Final artifacts, including the trained model and normalization parameters, were serialized using `final_model.save()` (H5 format) and `joblib.dump()` (GZ compression).

## Results

### Model Performance and Comparative Benchmarking

#### Validation against statistical noise

To assess the ability of StrucNS to capture genuine biophysical signals rather than statistical artifacts of amino acid composition, we compared its stability predictions against a noise model. We generated this control by randomly permuting feature values across all variants, preserving the underlying feature distribution while scrambling the predictive signal. In absolute stability (Δ*G*) classification, the noise model failed to exceed an F1 score of 0.20, while StrucNS reached 0.69 (TS1), 0.47 (TS2), and 0.48 (TS3) (Fig. 2B). Divergence in recall was comparably stark as the noise model plateaued at ∼0.19, whereas StrucNS achieved robust recall (Mean TS1: 0.64; TS2: 0.40; TS3: 0.42; Fig. 2B). This performance gap demonstrates the efficacy of our network-based approach.

#### Performance against large-scale PLMs

We next evaluated StrucNS against prominent PLMs, including ESM-2 [21], ESM-1v [14], and ESM-IF1 [27] (Fig. 2C). ESM-2 is a general-purpose transformer; ESM-1v is optimized for zero-shot variant prediction; and ESM-IF1 is an inverse folding model conditioning sequences on structure. These models are trained on vast datasets, ESM-2 and 1v on up to 250 million sequences (UniRef) and ESM-IF1 on 12 million AlphaFold structures, with parameters ranging from 650 million to 15 billion. With only ∼ 520K sequences of training, StrucNS achieved superior GMean across all test sets, particularly in de novo folds (TS2) where StrucNS maintained a GMean of 0.55 while ESM-2 collapsed to 0.31. ESM-1v yielded the lowest performance (GMean *<* 0.07), supporting the hypothesis that sequence-based evolutionary likelihoods struggle when the wild-type lacks a rich evolutionary history. ESM-IF1 and ProteinMPNN[22] also trailed StrucNS, with ProteinMPNN dropping to 0.17 in TS2. Crucially, while TS2 and TS3 represent true out-of-distribution regimes for StrucNS, they are likely represented within the massive training corpora of the ESM suite and ProteinMPNN. Despite this data disadvantage and a significantly smaller parameter count, StrucNS demonstrates higher resilience in zero-shot prediction by identifying the broader topological relationships defining a stable fold. Details of ESM and ProteinMPNN model implementations are added in SI Appendix, S3.1, S3.2, S3.3, S3.4.

#### StrucNS vs ThermoMPNN-D

We compared our network topology approach to ThermoMPNN-D [25] (SI Appendix, S3.5), a graph neural network (GNN)-based architecture trained directly on the same MegaScale dataset used for StrucNS. For mutational impacts (ΔΔ*G*), StrucNS outperformed ThermoMPNN-D in both TS1 and TS2 (Fig. 3D). Notably, while TS2 and TS3 are truly out-of-distribution for StrucNS, these data points were part of the explicit training set and priors for ThermoMPNN-D. Despite this, StrucNS matches or exceeds the performance of specialized GNN-based architectures.

**Figure 3.**
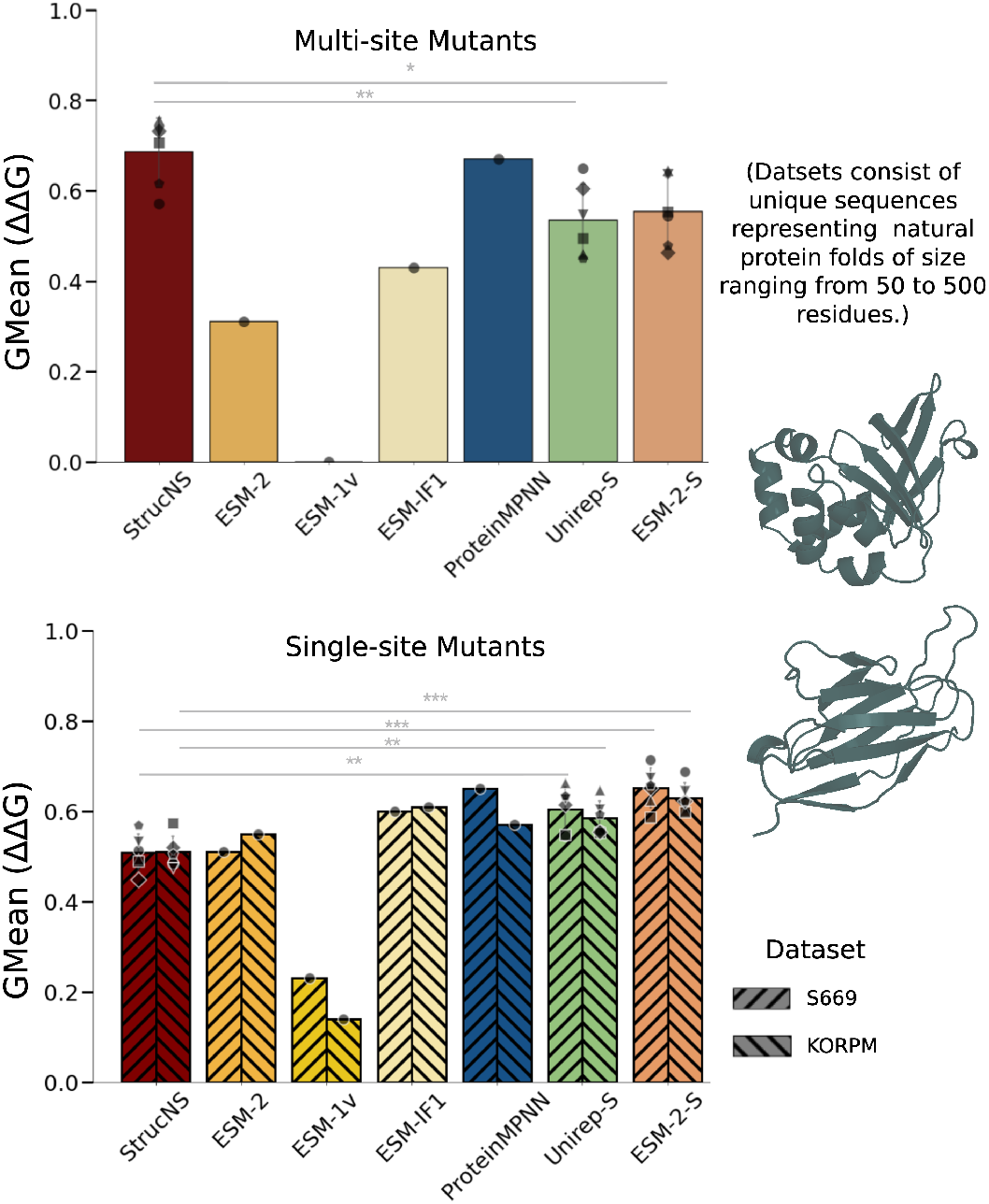
The predictive performance of StrucNS was evaluated in comparison to evolutionary models and high-capacity embeddings in PLMs. StrucNS was trained on miniproteins from Megascale dataset [26]. Data are presented for evaluations across multi-site mutants dataset (top) and single-site mutants dataset (bottom). Multi-site mutants dataset consists of curated data from literature having two or more mutations in proteins of size ranging from 50 to 500 residues and having large globular proteins. Single mutants dataset consists of datasets s669 and KORPM having single mutations and a size distribution of 40 to 600 residues. GMean for predicting mutational stabilization is shown.

#### Comparison with supervised PLMs

Finally, we compared StrucNS to supervised models utilizing high-capacity UniRep (SI Appendix, S3.7) and ESM-2 (SI Appendix, S3.6) embeddings (Fig. 3E). These supervised PLMs encode both evolutionary information and stability patterns through dual-round training in a black box manner, representing the performance ceiling. In the de novo regime (TS2), StrucNS (F1: 0.47) matched UniRep (F1: 0.48) but was less effective than the billion-parameter ESM-2 (F1: 0.64). On natural folds (TS3), StrucNS (F1: 0.39) was competitive, albeit less effective, versus UniRep (F1: 0.51) and ESM-2 (F1: 0.55). TS2 and TS3 are truly out-of-distribution for StrucNS but likely present in the PLM pre-training corpora; our method achieves reasonable performance using explicit structural connectivity rather than massive latent embeddings.

#### Prediction on single and multi-site mutations in large globular proteins

To evaluate the generalizability of StrucNS, we tested the model on standard single- and multi-site mutations of proteins that significantly exceed the size of the 40–72 amino acid miniproteins used during training. We utilized the PTMul [26, 48] dataset for multi-site mutations and the KORPM [26, 49] and s669 [50, 51] datasets for single-site mutations. PTMul contains multiple-site mutations derived from the ProTherm database, containing globular proteins featuring sizes typically ranging from 50 to over 500 residues. KORPM consists of single-point mutations curated from ProTherm and specifically curated to be balanced by removing an over-representation of alanine mutations and excluding data from extreme pH/temperature, covering a broad size distribution of 40 to 600 amino acids. The s669 dataset is a non-redundant set of mutations specifically filtered from ProTherm (non-overlapping with KORPM) and ensuring high-quality experimental ΔΔ*G* values, with protein lengths spanning 50 to 450 residues. Notably, these datasets are entirely out-of-distribution for StrucNS, whereas a significant portion of their data was included in the training sets of the comparative models. As illustrated in Fig. 3, StrucNS outperforms all other models in GMean when predicting the likelihood of mutational stability for multi-site mutants. This indicates that despite being trained on miniproteins, StrucNS captures generalized topological and structural patterns that govern stability across diverse folds. In single-site mutant predictions, StrucNS performs similarly to ESM-2 and ESM-IF1, while slightly underperforming compared to ProteinMPNN, Unirep-S, and ESM-2-S. These results highlight the robustness of StrucNS across varying sizes and physicochemistry. We hypothesize that StrucNS excels in multi-site mutant prediction because it utilizes network topology to predict absolute stability, allowing it to discern complex topological differences between multi-mutants and wild-type structures. ProteinMPNN, the other structure-based model analyzed, demonstrated performance similar to StrucNS in these evaluations. The lack of superiority of StrucNS in single-site mutants may result from reduced structural and connectivity differences in single mutants.

### Explainable AI reveals biophysical drivers of stability

Unlike black-box architectures, the StrucNS framework allows for a direct interpretation of the driving factors of protein stability. To deconstruct the biophysical drivers of stability, we utilized SHapley Additive exPlanations (SHAP) [52] to quantify the additive contribution of each feature to the model’s prediction (Fig. 4 and SI Appendix, Fig. S3). The primary driver, degree within intermediate subnetworks, shows that high connectivity within these hybrid polar/hydrophobic subnetworks correlates strongly with stability. The second most influential feature is inter-class fragment connectivity (ICFC) between intermediate and charged subnetworks, and the fifth is ICFC between intermediate and uncharged communities, while the fourth is low modularity in intermediate subnetworks. These four leading features indicate that the intermediate subnetwork must have high connectivity and be integrated across the entire protein network rather than being isolated within itself to ensure stability. Aligned with these leading features, importance share analysis reveals that the intermediate community features are the most influential for stability prediction, even more than hydrophobic communities (Fig. 4B). While the hydrophobic core is traditionally viewed as the primary determinant of stability, these unexpected results suggest that the highly connected intermediate region, bridging the polar surface and the hydrophobic core, exerts a more significant influence. Furthermore, the high inter-class fragment connectivity (ICFC) highlights the critical role of connectivity between functionally distinct protein regions in maintaining structural integrity.

**Figure 4.**
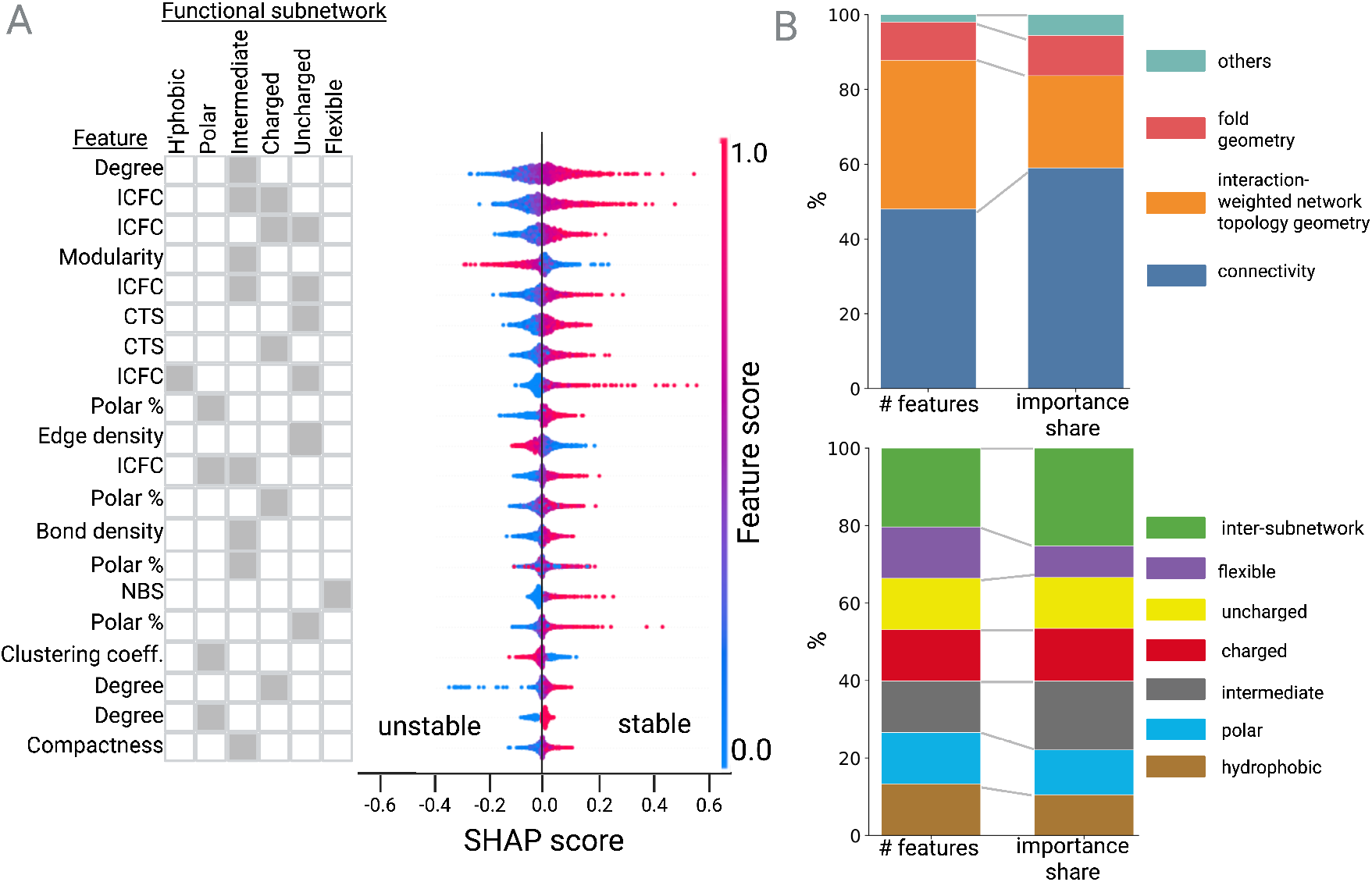
An explainable machine learning model reveals the topological determinants of protein stability. (A) *SHAP summary of biophysical drivers*. Individual topological features are ranked by their impact on model predictions using SHapley Additive exPlanations (SHAP). SHAP deconstructs a model’s decision-making by calculating the extent to which each feature shifts a specific prediction away from the baseline value (average model output across the training set; 0.25 likelihood of stability herein). Each point represents a protein variant. Its horizontal position on the x-axis indicates whether that feature’s value increased (positive SHAP) or decreased (negative SHAP) the prediction relative to the baseline. ICFC: inter-class fragment connectivity; CTS: class to community ratio. (B) (top) *Feature Importance Share*. The importance share of predictive signal is aggregated into four model stages: connectivity-based, geometry-based, projection-based, and others. (bottom) The number of features within each model stage is presented to enable comparison to the importance share.

More broadly, we computed the SHAP importance share for families of features as their fraction of the sum of absolute SHAP values across all features. Connectivity-based features – those computed purely from network – account for 59% of model importance, a significant rise from the 48% of such features (Fig. 4). Features that rely on the interaction-weighted network topology account for 28% of model impact. While influential, this share emerges from 117 features, 38% of all features; thus the relative impact per feature is much less than connectivity alone. Features computed from the original fold network topology contribute 10% from 10% of features. Details of implementation are provided in SI Appendix, S5.

### Model component analysis

#### Network topology is more significant than fold geometry

To further evaluate the contributions of network topology versus geometry, we systematically introduced noise into each component and measured the performance of the ablated models. Upon geometric noising (SI Appendix, S4.2) the ablated model maintained performance levels comparable to StrucNS across all test sets, as evidenced by the minimal impact on F1 and recall (Fig. 5). A second version of geometric noising yielded equivalent results (SI Appendix, S4.3). The results indicate that the precise spatial coordinates of the fold are minimally impactful to the model’s predictive power as compared to the influential connectivity and interaction-weighted geometry. We then evaluated the inverse condition by maintaining the protein geometry while randomizing the network topology (S4.1). By shuffling edges while preserving node degrees, we maintained network sparsity and connectivity but decoupled the topology from the physical fold. This manipulation led to a significant performance reduction across all metrics (Fig. 5). These results collectively demonstrate that network topology is a more potent predictor of protein stability than specific geometry. In this context, geometry serves primarily as a scaffold to establish the network topology or to recreate the latent physicochemical forces that drive protein folding and stability—a relationship that StrucNS successfully captures. Moreover, ablation of geometry in addition to ablating topology nominally but not substantially reduced performance relative to abalated topology alone (Fig. 5).

**Figure 5.**
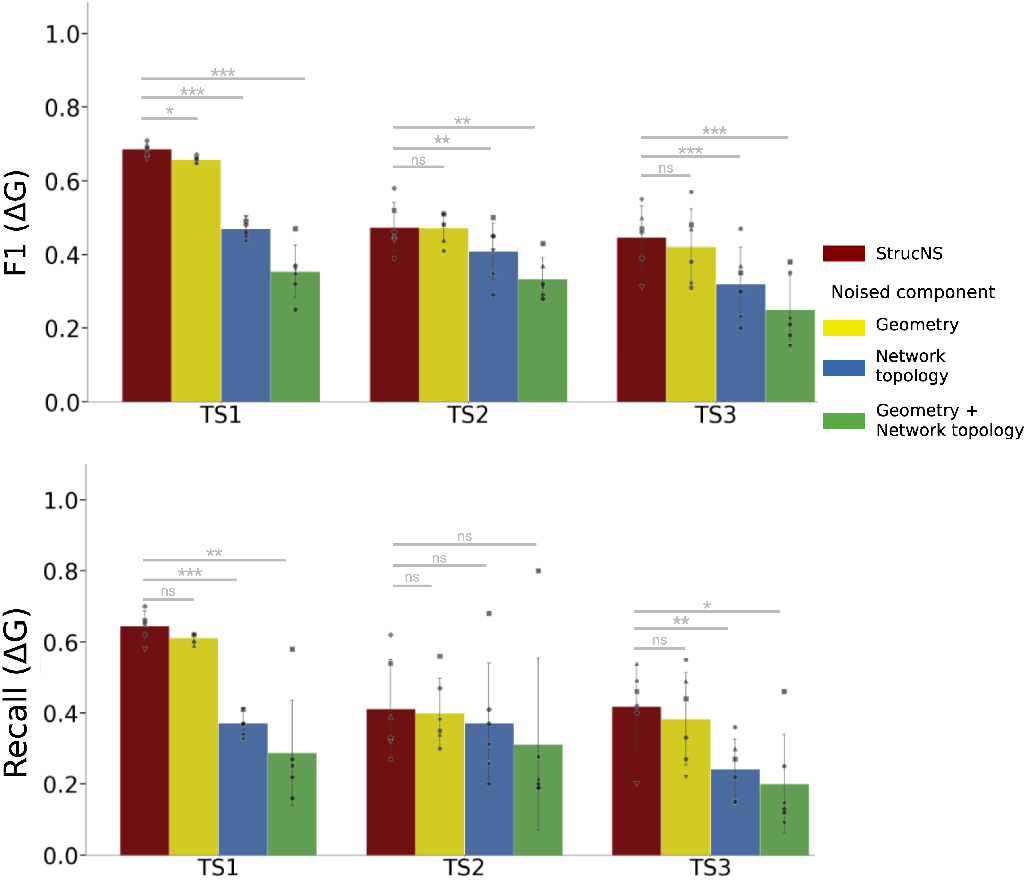
Systematic ablation of geometry and network topology reveals that network topology is the primary locus of the stability signal. The F1 (top) and Recall statistic (bottom) for predicting the stability classification (Δ*G >* 3 kcal/mol) of variants are shown for StrucNS and comparative models with model elements ablated. Geometry indicates randomized three-dimensional node positions in the protein fold. Network topology indicates the connectivity map comprising nodes, edges, and edge weights without any spatial information regarding the nodes. Geometry was noised using a random walk approach (yellow). Network topology was noised by randomizing edges in the network while conserving network sparsity and ensuring all nodes remained connected. Both geometry and network topology were randomized together (dark green and light green) to ablate the effects of geometry and network topology simultaneously. In these two cases, geometry was randomized using random box (dark green) and random walk (light green) approaches to ensure results were not an artifact of the randomization scheme. Finally, geometry was noised while peptide edges were maintained to reserve primary sequence information but exclude all non-peptide topology information.

#### Mechanistic necessity of functional subnetwork assignment

To quantify the predictive contribution of the functional subnetwork assignments, we randomly assigned communities within the interaction-weighted network projection and evaluated model performance, This ablation of physicochemical assignments significantly reduced F1 and recall (Fig. 6). Most strikingly, performance on TS2, the de novo protein folds, was reduced to the level of noise (Fig. 2). Thus, physicochemical-based subnetwork assignments provide critical utility to the StrucNS approach.

**Figure 6.**
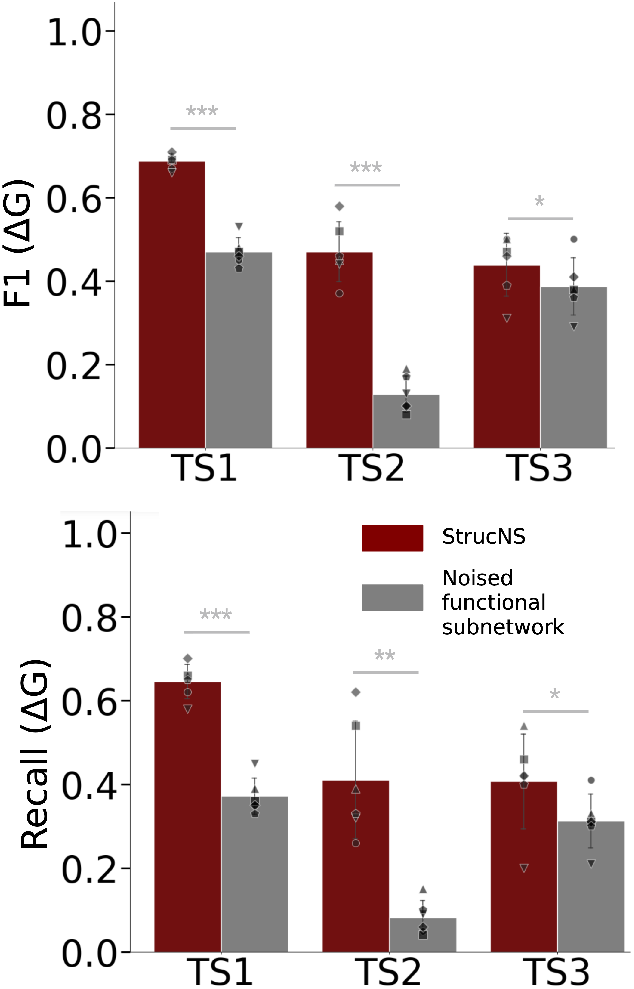
Ablation of functional subnetworks. The Recall (top) and F1 statistic (bottom) for predicting stability classification (Δ*G >* 3 kcal/mol) are shown for StrucNS and a comparative model with specific elements ablated. Functional subnetworks are ablated by randomly assigning detected communities to subnetworks, rather than labeling them by function. Specifically, a community is randomly assigned to random subnetwork A, B, or C. It is then further assigned to either random subnetwork D or E, and finally, potentially randomized as part of random subnetwork F. These assignments are performed while keeping the total number of subnetworks conserved.

#### Subnetwork-level vs. inter-subnetwork features

To resolve the multi-scale organization of the stability signal, we performed a hierarchical decomposition of the feature importance, categorizing SHAP contributions into subnetwork-level (aggregated community properties) and inter-subnetwork (pairwise interactions between functional modules) signals. Our results demonstrate that while the majority of the cumulative predictive power is stored in the aggregated properties of specific functional classes, the individual pairwise interactions exhibit a significantly higher average potency per feature (Fig. 4B). Analysis of the global importance share reveals that subnetwork-level features account for 75% of the total SHAP impact. Within this category, the model prioritizes the topological integrity of intermediate communities, as evidenced by the aforementioned dominance of degree and modularity in intermediate communities. Furthermore, the systematic appearance of percent polar across multiple subnetwork in the most influential features indicates that the model utilizes surface-core polarity distributions as a primary filter for identifying stably folded states. In contrast, inter-subnetwork (pairwise) features account for 25% of the importance share. This implies that the interaction between functional modules is also highly informative in the model. The prominence of ICFC metrics suggests that stability is sensitive to the specific “wiring” of the protein’s functional interface. These findings verify that the StrucNS framework operates through a dual-scale biophysical logic. The subnetwork-level features inform the macroscopic thermodynamic stability. Simultaneously, inter-subnetwork features capture the cooperative energy-coupling between distinct functional compartments.

#### Impact of peptide and non-peptide interactions

To evaluate the specific contribution of primary sequence connectivity versus higher-order structural interactions, we conducted an ablation study focusing on the edge types within the RIN, illustrated in Fig. 7. By selectively removing either peptide (backbone) edges or non-peptide interactions, we quantify how the model’s prediction signal depends on interactions. Performance metrics across all test sets indicate that non-peptide interactions are the primary drivers of high-fidelity stability prediction. In TS1 and TS3, the removal of non-peptide edges results in a significant decline in the F1-score and recall. Conversely, the removal of peptide edges results in a negligible performance change in TS1, with F1-scores remaining stable at ∼0.68 as well as less significant drop in TS3 relative to non-peptide edge removal. This suggests that the model’s topological features are robust enough to infer the linear backbone path implicitly from the density of non-covalent clusters. Interestingly, in TS2, the performance of the model without peptide bond edges remains equivalent to the base model.

**Figure 7.**
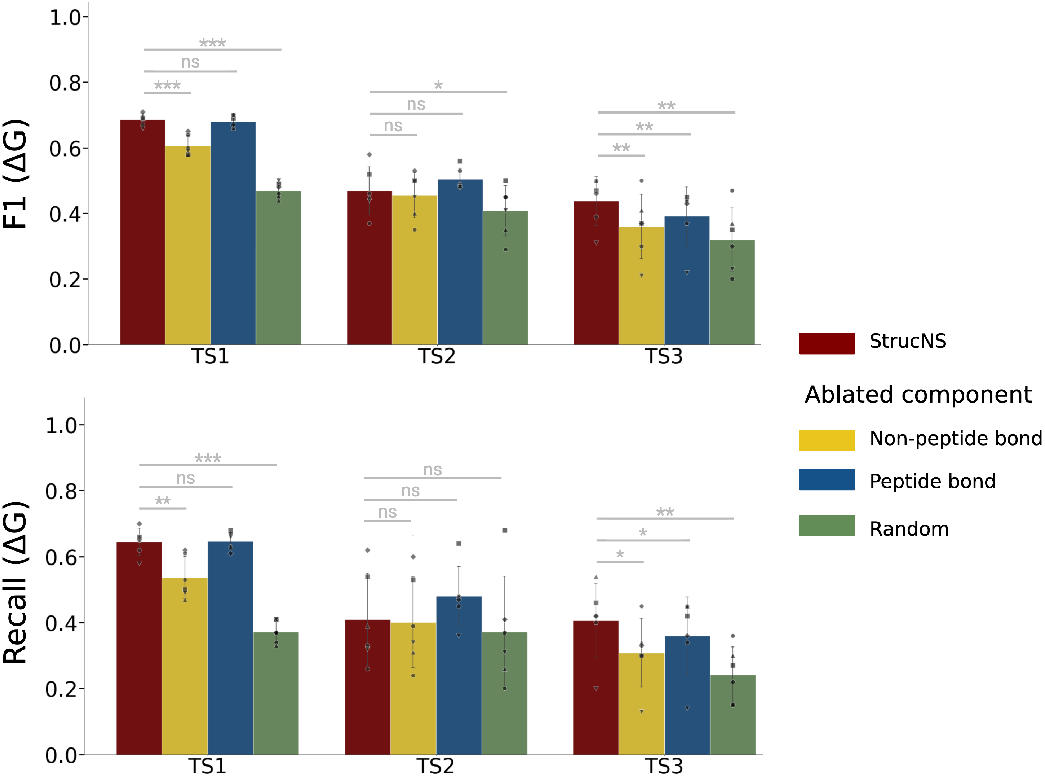
Systematic ablation of interaction types. The F1 (top) and Recall (bottom) for predicting stability classification (Δ*G >* 3 kcal/mol) are shown for StrucNS and comparative models with specific elements ablated. The network topology comprises peptide edges and various non-peptide edges. Peptide edges represent the protein backbone chain, while non-peptide edges define the tertiary protein fold. Non-peptide edges were removed (yellow) to understand the importance of tertiary fold without backbone chain information. Conversely, peptide edges were removed (blue) to evaluate the backbone chain’s contribution. Finally, all interaction types were randomized (green) to provide a benchmark for noise.

The critical nature of non-covalent wiring is further highlighted in the distribution of metrics for TS2 and TS3. In TS2, removing these edges causes the recall to remain low (∼0.24–0.39), while in TS3, the ablated *non-peptide bond* regime results in a precipitous drop in recall for specific cases, significantly underperforming the base model. This indicates that for more challenging or “noisy” structural datasets, the model relies almost exclusively on non-covalent topological invariants to distinguish stable folds from decoys. Without these edges, the model lacks the necessary “cross-talk” features—such as ICFC and inter-modular edge density—required to maintain predictive equilibrium in the absence of a perfect crystal structure.

## Discussion

In this study, we present StrucNS, a mathematical framework grounded in network theory that models protein fitness via interaction-weighted topology of the protein fold. While contemporary protein stability modeling is increasingly dominated by “black-box” architectures reliant on high-dimensional evolutionary embeddings, StrucNS offers a physically transparent alternative that derives its predictive signal directly from the explicit features of interaction-weighted network topology. By conceptualizing thermodynamic stability as an emergent property of network topology rather than a simple consequence of protein fold, we demonstrate that defined topological metrics serve as robust surrogates for the complex physicochemical forces governing the fitness landscape.

StrucNS, efficiently trained on ∼500,000 datapoints, achieves competitive, and in many cases superior, performance compared to state-of-the-art PLMs. Notably, in the *de novo* regime where evolutionary history is absent, StrucNS maintained a GMean of 0.55 while models like ESM-2 collapsed to 0.31 (Fig. 2), highlighting the resilience of a topology-based approach when navigating “fitness deserts”. Moreover, StrucNS was superior in predicting the likelihood of high stability in multi-mutants across a broad set of protein folds and sizes outside of the training regime (Fig. 3). Beyond mere prediction, the interpretability of StrucNS provides mechanistic insights into the structural logic of stability. SHAP analysis identifies the high connectivity and low modularity of intermediate (polar/hydrophobic mixed) subnetworks as primary drivers of stability. This suggests that the integration of the hydrophobic core with the polar surface through a highly connected intermediate region is more critical for structural integrity than the hydrophobic core alone.

Beyond elucidating fundamental insight of factors that dictate protein stability, StrucNS could empower protein stability engineering. Protein variants – identified via various search strategies (e.g., saturation mutagenesis, structure-guided mutagenesis, Bayesian optimization) – could be scored via StrucNS to identify mutants with increased likelihoods of high stability. In protein discovery workflows, StrucNS could be employed to evaluate the stability of a designed library or aid in scoring subsets of sequence space before proceeding to experimental validation. The mechanistic insight of StrucNS could also be leveraged to guide stabilization. StrucNS could be applied to identify the specific underperforming topological features that drove the designs toward instability. This feedback loop enables researchers to rationally identify residues for diversification within specific subnetworks, effectively shortening the design cycle. Future work can evaluate the efficacy of such techniques.

StrucNS benefits from the deep, high-integrity Megascale dataset of Tsuboyama et al.[26], which includes a broad set of protein folds and sequences, albeit focused on relatively small proteins (40 - 72 amino acids). Yet, the current implementation of StrucNS outperforms more broadly trained protein language models including on multi-mutants of a broad size range (Fig. 3). As additional datasets emerge, updated training can be performed to evaluate mechanistic insights – as well as predictive performance – across an even more expansive set of proteins. More broadly, the StrucNS network topology approach can be applied to other modes of protein performance including binding or additional metrics of developability.

The current performance and insight from StrucNS emerged from thoughtful design of model architecture and components. Despite the advancements, several avenues for future refinement remain. We utilized a classification approach, outputting the likelihood of stability being greater than 3 kcal/mol, to mitigate potential overfitting on high-throughput datasets with experimental variability. A stricter threshold was maintained to ensure a robust stable class. Future iterations may explore regression models as larger datasets become available. While feature analysis and ablation analysis revealed the relative impact of numerous design choices, further optimization can also be performed. Low-impact features, identified via SHAP analysis (Fig. 4), could be removed to enable more efficient training of impactful features. Interaction identification (distance thresholds and residue identities) and strengths can be tuned, and additional interaction types could be evaluated. Additional community classifications could also be tested. By integrating topological perspectives with emerging PLMs, one could develop synergistic models that combine the breadth of evolutionary grammar with the precision of structural network science toward a map of the protein fitness landscape.

## Author contributions statement and conflicts of interest

A.M., P.D., and B.J.H. designed research. A.M. performed research. A.M., P.D., and B.J.H. analyzed data. A.M., P.D., and B.J.H. wrote the paper. The authors declare that they have no competing interests.

## Funding and acknowledgments

We appreciate support from the NIH (R01GM146372 to B.J.H.). We appreciate assistance and computational resources from the Minnesota Supercomputing Institute.

## Supplementary Information

### StrucNS details

#### Network Construction and Interaction Criteria

The Residue Interaction Network (RIN) was generated using a custom Python pipeline incorporating the Biopython PDB parser and NetworkX. For all protein variants in our mutational libraries, 3D structural coordinates were generated via OmegaFold [42], as these high-fidelity predictions provide a consistent geometric basis for evaluating mutations in the absence of experimental structures. Residues were represented as nodes located at the centroid of all non-hydrogen atoms.

Edges were classified into six types based on distance cutoffs and atomic contact criteria. To ensure biophysical accuracy, aromatic interactions were identified using a centroid-to-centroid distance cutoff of 4.5 Å between aromatic rings (PHE, TRP, HIS, TYR), serving as a geometric proxy for orientations that enable *π*-*π* stacking. Further specific criteria for each interaction type are detailed below:

- **Peptide Bonds:** Defined between residues *i* and *i* + 1 along the polypeptide chain.
- **Hydrophobic Interactions:** Established between side-chain atoms of non-polar residues using a centroid-to-centroid distance cutoff of 5.0 Å.
- **Hydrogen Bonds:** Identified between donor and acceptor atoms with an atom-to-atom distance *<* 3.5 Å (4.0 Å for sulfur-containing donors).
- **Disulfide Bridges:** Defined between SG atoms of CYS residues within a 2.2 Å threshold.
- **van der Waals Forces:** Modeled as non-specific steric contacts for all residue pairs with a centroid distance *<* 5.0 Å not meeting the above specific criteria.

#### Weighting Schema and Energetic Justification

To model the relative rigidity and thermodynamic contribution of the protein fold, weights (*w*) were assigned based on representative bond dissociation energies (Δ*H*_*f*_) and empirical stability scales. The high weight assigned to peptide bonds (*w* = 10) enforces the primary backbone constraint as the most rigid component of the network. Non-covalent interactions were weighted to reflect their relative contributions to fold stability: hydrophobic contacts (*w* = 0.55) represent entropic core stabilization; hydrogen bonds (*w* = 0.28) represent polar stabilization; disulfide bridges (*w* = 1.0) account for covalent cross-links; aromatic stacking (*w* = 0.2) and van der Waals forces (*w* = 0.1) represent weaker, orientation-dependent or non-specific steric effects.

#### Community Detection (CD) via Modularity Maximization

The structural network was partitioned into communities using the Louvain heuristic, a two-phase iterative algorithm designed to maximize the modularity index (*Q*). In this framework, modularity serves as an objective function that quantifies the quality of a particular division of the network into communities. For a weighted graph, *Q* is defined as the difference between the actual weight of edges within communities and the expected weight if edges were placed at random while preserving the degree sequence of the nodes (the “background connectivity” null model). Mathematically, for a weighted network, modularity is expressed as:

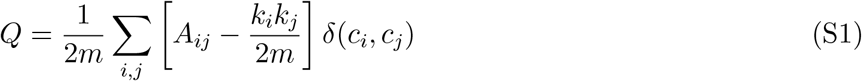

Where:

- *A*_i*j*_ is the weight of the edge between residue nodes *i* and *j*.
- *k*_*i*_ = Σ_*j*_ *A*_*ij*_ is the sum if the weights of the edges attached to node *i* (weighted degree).
- 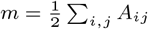 is the total sum of all edge weights in the network.
- *c*_*i*_ and *c*_*j*_ represent the community assignments of nodes *i* and *j*, respectively.
- *δ*(*c*_*i*_, *c*_*j*_) is the Kronecker delta function, which equals 1 if *c*_*i*_ = *c*_*j*_ and 0 otherwise.

The algorithm iteratively optimizes *Q* through two repeating phases: (i) local moving of nodes between communities to maximize local modularity gain, and (ii) aggregation of nodes assigned to the same community into a single “super-node” to form a new network for the subsequent pass. This variable assignment process continues until no further increase in *Q* is possible. This naturally discovers the optimal number of communities (*K*) for a given protein of *N* residues, where the condition *K < N* facilitates the transition from individual atomic centers to higher-order structural modules without the premature loss of structural information.

#### Hierarchical Spring-Balance Layout

To preserve the relative spatial orientation of structural modules while emphasizing interaction density, we implemented a hierarchical spring-balance layout using a modified Fruchterman-Reingold algorithm. The process is divided into two discrete tiers of optimization to reach a global topological equilibrium:

1. **Tier 1: Global Community Positioning**. A “community graph” is constructed where each node represents a coarse-grained community (*K*). Edge weights between community nodes *i* and *j* are calculated as the sum of all inter-community bond weights: 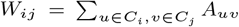. The global positions of these modules are determined by a 3D spring layout. By using the original PDB centroids as the initial state for the simulation, we maintain the native global topology while allowing the modules to shift toward a state of balanced tension.
2. **Tier 2: Local Residue Relaxation**. For each community *C*_*k*_, the internal residues are positioned relative to one another using a local spring-layout simulation. The internal edge weights *A*_*uv*_ (where *u, v* ∈ *C*_*k*_) define the attractive spring constants. Once local equilibrium is reached, the residues are translated such that their local centroid aligns with the global community position calculated in Tier 1.

The force *F* acting on a node in this system is defined by the summation of attractive forces *f*_*a*_ (along edges) and repulsive forces *f*_*r*_ (between all node pairs):

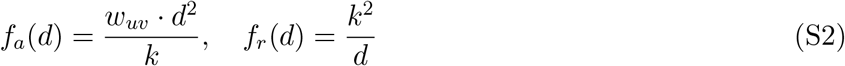

where *d* is the distance between nodes, *w*_*uv*_ is the physicochemical bond weight, and *k* is the optimal distance parameter derived from the expected node density. This hierarchical approach ensures that the resulting “topological coordinates” reflect the physical equilibrium of the interaction network. By allowing nodes to migrate under these forces, the layout acts as a noise-reduction filter; residues that appear buried in a noisy structure but lack strong stabilizing connectivity will drift toward the periphery, while those forming the true structural core will be pulled closer together.

#### Multi-Label Functional Classification Logic

To translate topological communities into biologically interpretable modules, we categorized each community *C* based on the intrinsic properties of its constituent residues. This classification relies on two primary biophysical scales: the Kyte-Doolittle scale, which measures desolvation energy, and the Grantham scale, which accounts for side-chain polarity and volume.

For each community, a Hydrophobicity Score (*H*_*C*_) was calculated as the arithmetic mean of the normalized scores of its residues. To ensure the classification was robust against structural noise, we applied a three-tiered labeling logic:

1. **Hydrophobicity Regime:** Each community is assigned exactly one label based on its mean score *H*_*C*_ :
  - **Hydrophobic:** *H*_*C*_ ≥ 0.66, identifying dominantly non-polar core fragments.
  - **Polar:** *H*_*C*_ ≤ 0.33, identifying dominantly hydrophilic surface fragments.
  - **Mixed:** 0.33 *< H*_*C*_ *<* 0.66, identifying amphipathic or transition regions.
2. **Charge Status:** Each community is assigned exactly one label based on the fraction of ionizable residues (*f*_*charge*_):
  - **Charged:** If *f*_*charge*_ *>* 0.3 (where residues are Arg, Lys, Asp, or Glu).
  - **Uncharged:** If *f*_*charge*_ ≤ 0.3.
3. **Flexibility Propensity:** A community is tagged as **Flexible** if the fraction of residues associated with high conformational entropy (*f*_*flex*_, specifically Gly and Pro) exceeds 30% (*f*_*flex*_ *>* 0.3).

This framework ensures that every module is described across three orthogonal biophysical axes. For example, a module might be classified as “Hydrophobic-Uncharged” while another is “Polar-Charged-Flexible.” This nuanced characterization is necessary to preserve the subtle biophysical nuances of the protein fold during the subsequent feature aggregation steps, where topological invariants are mapped to these functional categories.

#### Hierarchical Aggregation Logic

The transformation from raw residue data to the final machine-learning-ready feature vector follows a two-step hierarchical protocol.

##### Step 1: Node-to-Community Aggregation

For a community *C* containing a set of nodes *V*_*C*_, the community-level feature *f*_*C*_ is calculated as:

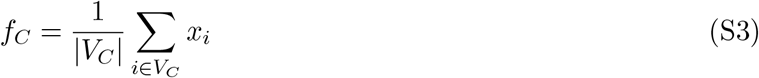

##### Step 2: Community-to-Class Aggregation

Communities are grouped by physicochemical labels. We derive class-level features *F*_*Class*_ via three strategies:

- **Node-Weight Mapping (NWM):** 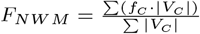
- **Volume-Weight Mapping (VWM):** 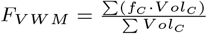
- **Simple Mean (SM):** 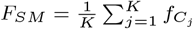

#### Comprehensive Feature Definitions

To ensure reproducibility and clarity, this section provides the formal mathematical definitions of the features extracted from the residue interaction networks. These features are organized into four functional categories and aggregated using three distinct strategies: **Node-Weighted Mean (NWM), Volume-Weighted Mean (VWM)**, and **Simple Mean (SM)**.

#### Aggregation Strategies

The community-level metrics (*F*_*k*_) are aggregated into class-level descriptors (*F*_*C*_) as follows:

- **Node-Weighted Mean (NWM)**: 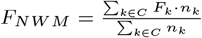, where *n*_*k*_ is the number of residues in community *k*.
- **Volume-Weighted Mean (VWM)**: 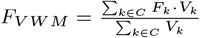, where *V*_*k*_ is the spring-model convex hull volume.
- **Simple Mean (SM)**: 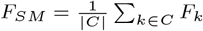, representing the unweighted average across all communities in class *C*.

#### 1. Connectivity Based Features

These metrics quantify the topological robustness and communication efficiency of the structural subnetworks.

- **Weighted Degree (***D*): The average sum of edge weights *w*_*ij*_ incident to residues within the community.
- **Clustering Coefficient (***CC*): Measures the local transitivity within the community graph.
- **Centrality Metrics**: Includes *Betweenness Centrality* (shortest-path control), *Eigenvector Centrality* (influence), and *PageRank* (importance based on neighbor quality).
- **K-Core Decomposition**: The level of the maximal subgraph where all residues have at least *k* neighbors.
- **Modularity (***M* ): The degree to which the community structure is optimized relative to a null model.
- **Edge/Bond Density**: *Edge Density* is the ratio of existing to potential edges; *Bond Density* is the average weight per edge: 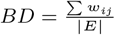.
- **Algebraic Connectivity (**λ_2_): The second smallest eigenvalue of the Laplacian matrix, indicating subnetwork synchronization and stability.
- **Topological Density Factor (TDF)**: A hybrid metric defined as the ratio of topological edge density to the physical convex hull volume: 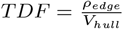.
- **Inter-Class Boundary Fraction (ICBF)**: Ratio of the sum of weights for edges connecting Class A and B to the total weighted degree (perimeter) of the boundary nodes.
- **Class Territorial Overlap (CTO)**: The volumetric intersection ratio between two class territories: 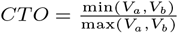.
- **Inter-Class Force Coefficient (ICFC)**: The total interaction weight between classes normalized by the maximum individual bond weight observed.

#### 2. Geometry Based Features

Metrics focused on the physical distances and spatial resolution of the residue clusters.

- **Average Pairwise Distance (APD)**: The mean Euclidean distance between all possible pairs of residues in the community.
- **Network Resolution Factor (NRF)**: The ratio of the spatial spread in PDB coordinates (*σ* _*PD B*_) to the spread in spring-relaxed coordinates 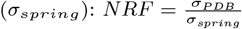 .
- **Curvature Correction Ratio (CCR)**: The ratio of compactness between the native and equilibrated states: 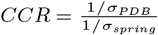.
- **Centroid-Induced Coordinate Shift (CICS)**: The Euclidean distance between the community’s PDB-centroid and its spring-model centroid.
- **Class Topological Stability (CTS)**: The ratio of inter-community weights to intra-community weights within a functional class.

#### 3. Projection Based Features

These features project structural data into 3D spatial descriptors to characterize the global protein envelope.

- **Convex Hull Volume (***V*_*ch*_): The volume of the smallest convex polyhedron containing all residue coordinates in the community.
- **Spread (***σ*): The standard deviation of residue distances from the community centroid.
- **Proximity Centralization**: A measure of subnetwork density relative to its geometric center: 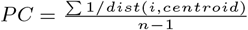.
- **Structural Tightness Coefficient (STC)**: The fractional volume occupied by a class relative to the total volume of the complex.
- **Z-Centroid (***Z*_*cent*_): The distance from the community centroid to the global center of mass of the protein.
- **Centroid Coordinates (X, Y, Z)**: The mean spatial projection of the class in 3D space.
- **Weighted Inter-Class Distance Spread (WICDS)**: The standard deviation of all pairwise Euclidean distances between residues of two different functional classes.

#### 4. Chemical and Functional Features

- **Polar Percentage (***PolarPct*): The fraction of residues in the community belonging to polar or charged categories (GLU, ASP, LYS, ARG, HIS).

## Additional results

### Distribution of Functional Subnetworks vs. SASA

To characterize the structural environment of the predicted functional subnetworks, we quantified the solvent accessibility of each constituent residue across the three physicochemical classes. Solvent Accessible Surface Area (SASA) was calculated using the Shrake-Rupley algorithm as implemented in the Biopython structural bioinformatics package. To account for the inherent size differences among amino acid side chains, raw SASA values were converted to Relative Solvent Accessibility (RSA) by normalizing against the maximum possible SASA (*S*_*max*_) for each residue type in a Gly-X-Gly tripeptide context, using reference values from Miller et al. (1987). This normalization yields a scale where 0.0 represents a fully buried residue and 1.0 represents maximum solvent exposure.

The resulting structural consensus, derived from *n* = 17, 135 protein environments, reveals distinct spatial partitioning (Fig. S1). The *Hydrophobic* subnetwork exhibits a sharp density peak at low RSA values near 0.2, confirming that these predicted functional nodes predominantly localize within the protein core to drive stability via the hydrophobic effect. Conversely, the *Polar* subnetwork displays a significant rightward shift with a density maximum at approximately 0.6 RSA, consistent with the requirement for polar residues to reside on the protein surface to maintain solubility and facilitate hydrogen-bonding with the aqueous solvent. The *Mixed* subnetwork serves as a structural intermediate, with a broad distribution centered near 0.55, suggesting that these residues form a transitional interface between the hydrophobic interior and the solvent-exposed exterior. The relationship between the localized residue class and its solvent environment is captured by the mean Kernel Density Estimate (KDE) for each class *c*, calculated as:

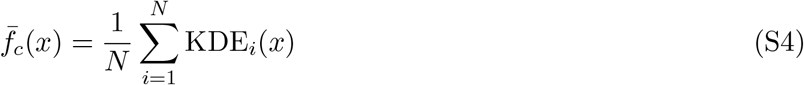

**Figure S1.**
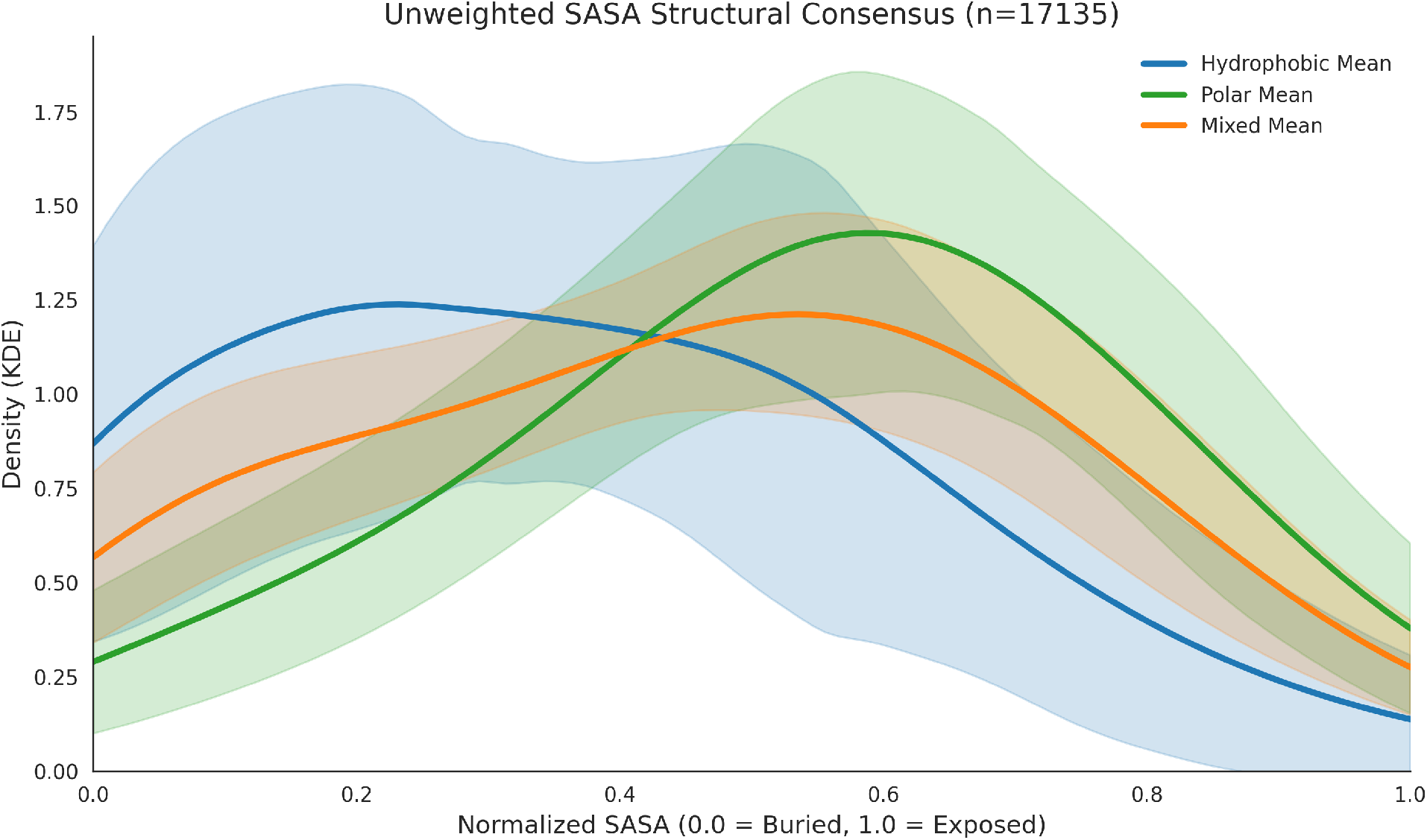
Structural consensus of SASA distributions for predicted functional subnetworks. The plot illustrates the unweighted mean KDE densities (solid lines) and standard deviations (shaded regions) for Hydrophobic (blue), Polar (green), and Mixed (orange) residue classes (*n* = 17, 135). The clear spatial segregation between the buried hydrophobic core and the exposed polar surface validates the model’s ability to discern structurally relevant residue environments.

**Figure S2.**
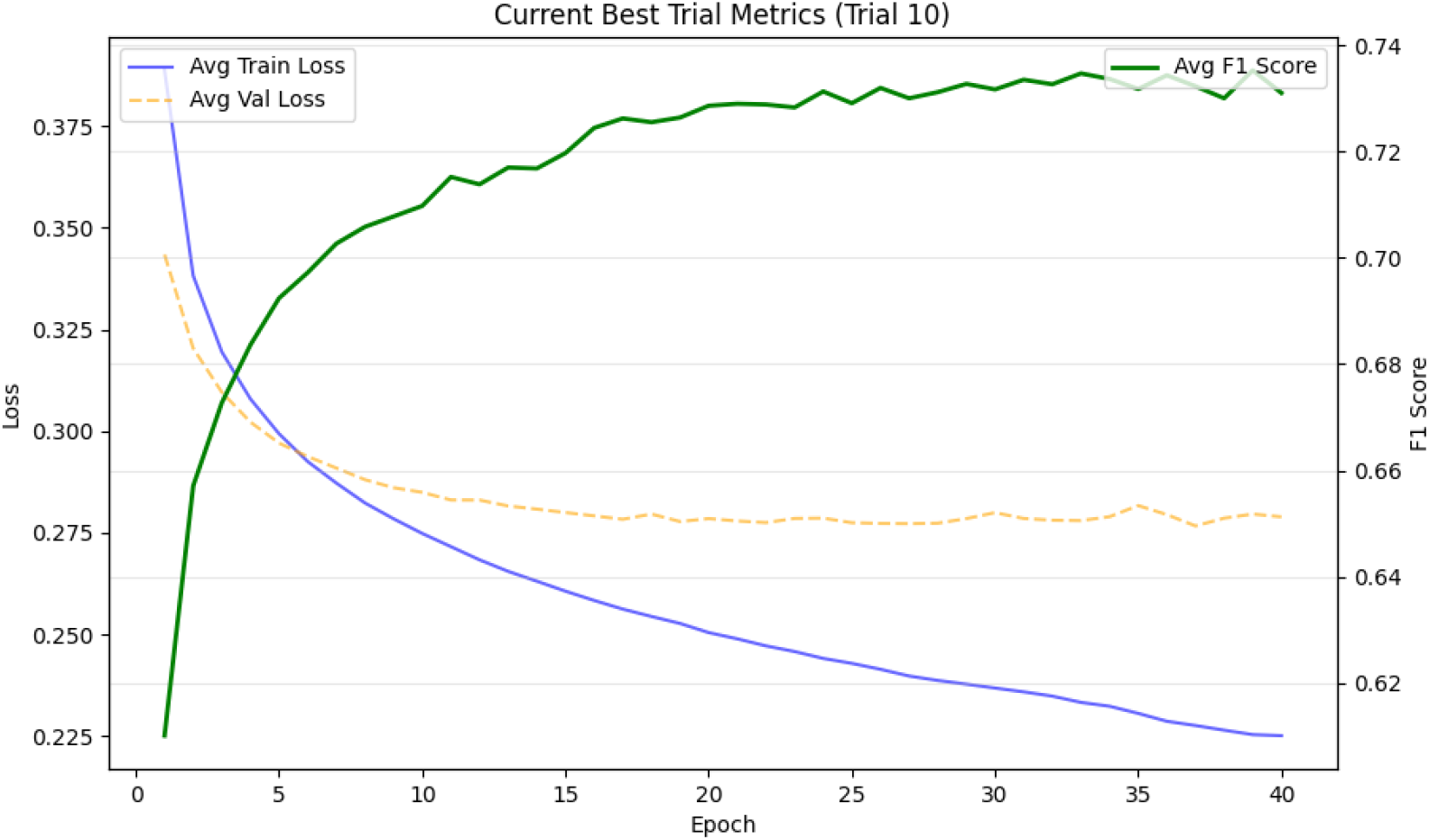
Training and validation metrics for the optimized Δ*G* prediction model. The primary y-axis (left) displays the binary cross-entropy loss for training (solid blue) and validation (dashed orange) sets. The secondary y-axis (right) denotes the *F*_1_ score (solid green) achieved across the validation folds. Convergence is reached near epoch 40, where the *F*_1_ score stabilizes at *≈* 0.73, indicating high model reliability in identifying stable variants.

where *x* represents the normalized RSA and *N* is the number of proteins. The observed robustness of these distributions demonstrates that the deep learning model’s predicted features are grounded in fundamental principles of protein architecture. These results provide biophysical validation that the functional subnetworks identified by the model correspond to specific, structurally conserved niches within the protein fold.

### Monitoring overfitting

To evaluate the stability and generalizability of the deep learning architecture, we monitored the training dynamics across the cross-validation folds. Figure S2 illustrates the evolution of the primary performance metrics for the optimal hyperparameter configuration for one of the six ensemble models of StrucNS. All of the six model ensembles followed the same pattern while monitoring for overfitting. The *Average Training Loss* (solid blue line) demonstrates a monotonic decay, asymptotically approaching a minimum value near 0.225, which signifies effective minimization of the objective function.

Parallel to the training trajectory, the *Average Validation Loss* (orange dashed line) exhibits a rapid initial descent before stabilizing at approximately epoch 20. The absence of a characteristic “U-shaped” divergence in the validation loss confirms that the model successfully mitigated overfitting, maintaining a minimal generalization gap. This stability is largely attributable to the implementation of family-aware data partitioning and the optimized dropout-regularization layers.

Classification performance was primarily assessed via the *Average F*_1_ *Score* (solid green line), which provides a balanced metric for identifying high-stability variants (Δ*G >* 3). As shown in the dual-axis plot, the *F*_1_ score experiences a sharp increase during the first 15 epochs, eventually plateauing at a robust value of approximately 0.73. The synchronization between the stabilization of the validation loss and the plateauing of the *F*_1_ score indicates a well-calibrated learning rate and effective feature extraction.

The optimization objective, defined as the *Mean Maximal* 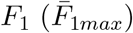, was calculated to ensure the final model captured the peak discriminatory power achieved during the training process:

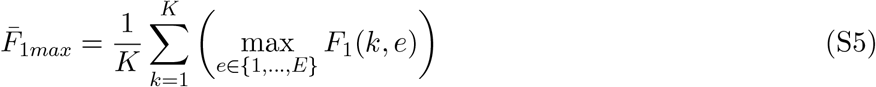

where *K* = 5 represents the cross-validation folds and *E* is the epoch count. To preserve the most generalizable parameter state, we utilized a weight-restoration protocol that isolated the specific epoch *e*^*^ corresponding to the global maximum of the validation *F*_1_ curve. This methodology ensures that the final predictive model represents the optimal compromise between sensitivity and specificity before the onset of marginal stochastic fluctuations observed in the later epochs (*e >* 35).

## Control model implementation

### ESM-2 Implementation and Evolutionary Scoring

The evolutionary fitness of protein variants was quantified using the Evolutionary Scale Modeling (ESM-2) transformer architecture. We employed the `esm2_t33_650M_UR50D` pre-trained model, consisting of 33 layers and 650 million parameters, implemented via the `esm` (v2.0.0) library in `PyTorch` (v2.0). All inference tasks were executed on NVIDIA A100 GPUs to accelerate the transformer attention mechanism.

### Sequence Extraction and Preprocessing

Primary sequences were derived from structural coordinates to ensure parity between structural network analysis and sequence-based scoring. Amino acid sequences were extracted from the ATOM records of PDB files using `Bio.SeqIO.parse` with the `“pdb-atom”` flavor. Sequences exceeding the 1022-residue context window were truncated to meet the transformer’s positional encoding constraints.

### Pseudo-Log-Likelihood (PLL) Objective

To evaluate sequence stability, we calculated the pseudo-log-likelihood (PLL) based on the Masked Language Modeling (MLM) objective. For a sequence *S*, the PLL was computed by iteratively applying the `alphabet.mask_idx` to each position *i* and summing the log-probabilities of the original amino acid *x*_*i*_ as predicted by the model’s logits:

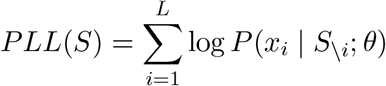

where *S*_\*i*_ represents the sequence with the *i*-th residue masked, and *θ* denotes the model parameters. The relative evolutionary impact of a mutation was determined by the Log-Likelihood Ratio (LLR):

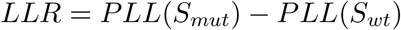

### Computational Optimization

The scoring pipeline was optimized for large-scale datasets using a dual-caching strategy. Calculated *PLL*(*S*_*wt*_) values were stored in-memory (`Dict[str, float]`) and persisted to disk via `csv.DictWriter` to eliminate redundant computations for common wild-type backgrounds. The workload was parallelized across high-performance computing clusters using SLURM job arrays, where the `SLURM_ARRAY_TASK_ID` environment variable was mapped to partition-specific CSV inputs for distributed processing.

### ESM-1v implementation and variant scoring

To establish a baseline for variant effect prediction using zero-shot evolutionary information, we implemented an ensemble of five `ESM-1v` transformer models (esm1v_t33_650_MUR90S_[1-5]) . Each model in the ensemble consists of 33 layers and 650 million parameters, trained with different random seeds on the UniRef90 database to improve the robustness of the predicted mutational effects.

### Ensemble Model Loading and Architecture

The ensemble was instantiated using the `esm.model.esm1.ProteinBertModel` class. Because the models were loaded from raw state dictionaries, we implemented a custom `Namespace` to define the `roberta_large` architecture parameters: 33 layers, an embedding dimension of 1280, 20 attention heads, and a feed-forward network `(ffn_embed _dim)` of 5120. A clean-up logic was applied to the `state _dict` keys to resolve naming conflicts across the five checkpoint iterations, and the language modeling heads `(lm_head)` were dynamically resized to match the target vocabulary.

### Wild-type Marginal Scoring Strategy

Variant scoring was executed using the “Wild-type Marginal” strategy, which is computationally efficient for single and multi-point mutations. For each variant, we identified the set of indices {*pos*} where the mutant sequence (*S*_*mut*_) differed from the wild-type (*S*_*wt*_). The Log-Likelihood Ratio (*LLR*) was computed by passing the wild-type sequence through the ensemble and comparing the log-probabilities of the mutant and wild-type amino acids at the affected positions:

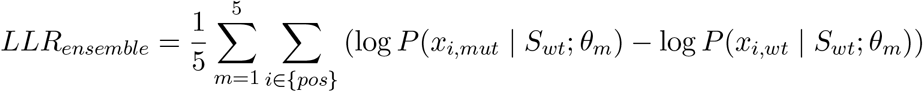

where *x*_*i,mut*_ and *x*_*i,wt*_ are the mutant and wild-type residues at position *i*, and *θ*_*m*_ represents the parameters of the *m*-th model in the ensemble. This approach captures the probability of the mutation within the context of the wild-type sequence.

### Implementation and Parallelization

Primary sequences were extracted from PDB ATOM records using `Bio.SeqIO` to ensure exact correspondence with the structural models. The inference pipeline was parallelized using SLURM job arrays, with each task processing a distinct partition of the dataset. To ensure reliability during long-running jobs, we implemented a file-stream logic using `csv.DictWriter` and `f.flush()`, allowing for real-time data persistence and the ability to skip previously processed variants upon job resumption.

### ESM-IF1 Structural Inverse Folding Implementation

To evaluate the structure-sequence compatibility of protein variants, we implemented the `ESM-IF1` inverse folding model `(esm_if1_gvp4_t16_142M _UR50)` . This model utilizes a Geometric Vector Perceptron (GVP) Graph Neural Network (GNN) combined with a transformer decoder (142 million parameters) to map 3D structural coordinates to sequence probabilities.

### Coordinate Loading and Feature Extraction

Structural inputs were sourced from OmegaFold-predicted PDB files. The model interface utilized the `esm.inverse_folding.util.load_cords` function to extract the backbone coordinates (*N, Cα, C*) of the protein chain. These coordinates serve as the invariant structural backbone upon which the likelihood of both wild-type and mutant sequences was assessed.

### Inverse Folding Scoring Objective

The model computes the conditional log-likelihood of a sequence given its 3D backbone coordinates. We quantified the structural stability of each variant by calculating the log-likelihood (*ℒ*) using the `score sequence` method. The fitness of a mutation was determined by the change in structural log-likelihood (Δ*ℒ*_*IF*_) between the mutant structure and the wild-type structure:

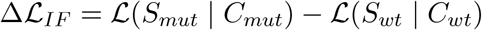

where *S* represents the amino acid sequence and *C* represents the 3D atomic coordinates. This metric captures the degree to which a specific mutation is favored or disfavored by the local and global structural environment.

### Computational Optimization and Scalability

The implementation featured several performance-enhancing strategies:

- **Dynamic Coordinate Mapping:** An enhanced search logic was implemented in `find_pdb_path` to resolve varied file naming conventions (e.g., `.pdb_wte.pdb` vs. `.pdb.pdb`) across heterogeneous PDB directories.
- **Persistence and Cache Recovery:** To minimize redundant GNN passes, wild-type structural scores were cached in a `wt_score_cache` (`Dict[str, float]`). Upon execution, the script automatically resumed by cross-referencing existing output files to reconstruct the cache and skip completed entries.
- **Inference Monitoring:** Real-time telemetry was implemented to track the average processing time per structural record, providing projected completion windows for large-scale SLURM job arrays.

### ProteinMPNN Structural Scoring and Evaluation

To assess the structural stability and energetic favorability of variants, we utilized `ProteinMPNN` (v.1.0.1), a graph-based neural network architecture designed for inverse folding. The model evaluates how well a specific sequence fits a given three-dimensional backbone by calculating the sequence log-likelihood based on structural node and edge features.

### Model Execution and Parameterization

The scoring pipeline was implemented via a `subprocess` wrapper in `Python`, calling the `protein_mpnn_run.py` script within a dedicated `Conda` environment. We employed the `v 48 020` model weights, which provide a balance between computational efficiency and accuracy. Scoring was restricted to Chain A `(--pdb_path_chains “A”)` with a `batch_size` of 1 to ensure deterministic score extraction for each specific wild-type and mutant PDB coordinate set.

### Log-Likelihood Extraction and Differential Scoring

The `ProteinMPNN` score, representing the average negative log-probability of the sequence given the backbone, was parsed from the resulting FASTA output using regular expression (`re`) matching:

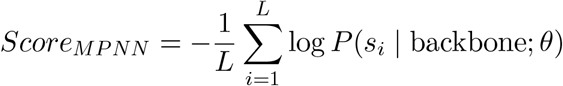

For each variant, we calculated the scores for both the wild-type (*LL*_*WT*_) and mutant (*LL*_*mut*_) structures. The structural impact was quantified as the Log-Likelihood Ratio (*LLR*_*MPNN*_):

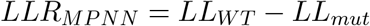

In this convention, a positive *LLR* indicates that the mutation is less compatible with its structure compared to the wild-type, suggesting a potential loss of stability or fitness.

### Automated Pipeline and Data Management

Computational throughput was optimized through several implementation strategies:

- **Dynamic Directory Mapping:** The script utilized a custom `get_mpnn_score` function to handle complex PDB naming conventions, automatically resolving suffixes (e.g., `.pdb_suffix.pdb`) associated with mutant structures.
- **Parallelized Batch Processing:** Distributed execution was managed via SLURM job arrays, where the `SLURM ARRAY TASK ID` was utilized to partition the input datasets into discrete tasks.
- **Resource Cleanup and Persistence:** To avoid filesystem overhead during large-scale scoring, a temporary working directory (`temp_mpnn_[PID]`) was created for each process to store intermediate FASTA outputs. This directory was purged using `shutil.rmtree` upon successful score extraction and row-wise persistence to the output CSV.

### ThermoMPNN Thermodynamic Stability Prediction

To provide explicit thermodynamic benchmarks, we implemented the `ThermoMPNN` model, which extends the ProteinMPNN architecture with a dedicated stability prediction head. We utilized the `single` configuration of the `v2` model, specifically parameterized to predict ΔΔ*G* values for single-point mutations based on 3D structural environments.

### Model Architecture and Featurization

The implementation utilized a hierarchical featurization process via `tied_featurize_mut` from the `thermompnn.datasets` module. Structural data were extracted from OmegaFold-generated PDB files and converted into graph-based features including backbone coordinates (*X*), amino acid sequences (*S*), and residue indices. To ensure numerical stability during the forward pass, we applied `torch.nan_to_num` to the coordinate tensors.

### Forward Pass and ΔΔG Mapping

Stability predictions were generated by passing the featurized structural graphs through the ProteinMPNN encoder. We extracted the hidden states `(all_mpnn_hid)` from the final model layers and applied a light attention mechanism `(model.light_attention)` to aggregate local structural contexts. The final ΔΔ*G* map was produced by the `model.ddg_out` head. To ensure the predictions were relative to the starting structure, we normalized the output map by subtracting the wild-type stability score:

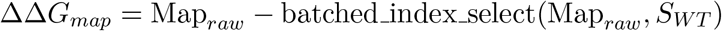

The final prediction for a specific mutation *X*_*i*_*Y* (where *X* is the wild-type residue at position *i* and *Y* is the mutant) was retrieved by indexing the normalized `ddg_map` at the corresponding position and amino acid identity.

### Automated Pipeline for Large-Scale Mutagenesis

The computational pipeline integrated several custom modules to handle high-throughput evaluation:

- **Mutation Parsing:** We developed a regex-based parser (`parse_mutation`) to extract wild-type identity, sequence position, and mutant identity directly from standardized GraphML filenames.
- **Priority-Based PDB Resolution:** The `get_wt_pdb_path` function implemented a multi-directory search logic to resolve wild-type structures across distinct processing tiers (e.g., `omegafold_pdbs_WT` and secondary search paths).
- **Fault-Tolerant Execution:** The script featured a robust error-handling framework that logged specific failure modes (e.g., `IndexError`, `ModelError`, or `WT_NotFound`) directly to the output CSV, ensuring dataset integrity even when structural data were missing or corrupted.

### Supervised ESM-2 Embedding and Classification

To leverage the high-level semantic information captured by large-scale protein language models, we implemented a supervised learning framework that combines fixed feature extraction from a pre-trained transformer with a downstream deep neural network optimized for stability classification.

### ESM-2 Feature Extraction and Mean Pooling

Sequence-level representations were generated using the `esm2_t33_650M_UR50D` architecture. For each sequence, hidden states were extracted from the final transformer layer (*layer* = 33), resulting in a 1280-dimensional vector per residue. To obtain a fixed-length global representation for each protein, we applied a mean-pooling operation across the sequence dimension (excluding `<cls>` and `<eos>` tokens):

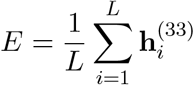

where *L* denotes the sequence length and 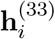 represents the hidden representation of residue *i*. Inference was performed on NVIDIA A100 GPUs using a checkpoint-aware script to ensure data persistence and reproducibility.

### Data Preprocessing and Dimensionality Reduction

Prior to training, the 1280-dimensional embeddings were processed through a multi-stage pipeline:

- **Imputation:** Missing values were addressed using a `SimpleImputer` with a mean-value strategy.
- **Standardization:** Features were normalized to zero mean and unit variance using `StandardScaler` .
- **Principal Component Analysis (PCA):** To mitigate the curse of dimensionality and prevent overfitting, we applied PCA, retaining enough components to explain 99% of the total variance.

### Hyperparameter Optimization and Model Architecture

The downstream classifier was developed using `TensorFlow/Keras` . The target variable was binarized based on experimental Δ*G* values from the Tsuboyama et al. dataset, defining stable variants (*y* = 1) as those with Δ*G >* 3 kcal/mol.

To determine the optimal architecture, we utilized `Optuna` for 300 trials of Bayesian optimization. The search space included the number of hidden layers (1–4), layer widths (32–512 units), dropout rates (0.1–0.5), and the conditional application of Batch Normalization. A family-aware splitting strategy was employed to ensure that proteins from the same evolutionary family were restricted to either the training or validation sets, preventing data leakage.

### Training and Evaluation

The final model was retrained using the best hyperparameters (`Adam` optimizer, variable learning rates). We implemented a custom `F1ScoreCallback` to monitor the F1-score at each epoch and restore weights associated with the highest validation performance:

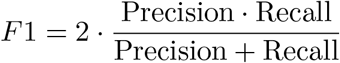

The model was evaluated across three independent test sets, with performance logged via confusion matrices and class-specific recall to assess the model’s robustness across diverse protein scaffolds.

### Supervised UniRep Embedding and Classification Pipeline

To capture the statistical and structural features encoded within protein primary sequences, we implemented a supervised framework utilizing UniRep—a multiplicative Long Short-Term Memory (mLSTM) recurrent neural network—to generate fixed-length vector representations for stability classification.

### UniRep Feature Extraction and mLSTM Processing

The UniRep embeddings were generated using the `jax_unirep` framework. Sequences were standardized and filtered to include the 20 standard amino acids and the unknown residue token (`X`). The mLSTM architecture processed each sequence to produce 1900-dimensional hidden states. To obtain a global representation for each variant, a mean-pooling operation was applied:

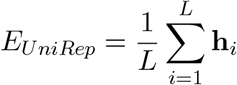

where *L* is the sequence length and **h**_*i*_∈ ℝ 1900. Computational throughput was optimized by parallelizing extraction across high-performance computing clusters using Slurm job arrays to partition the dataset into discrete chunks.

### Preprocessing and Dimensionality Reduction

Prior to classification, the 1900-dimensional vectors underwent a robust preprocessing pipeline:

- **Family-Aware Stratification:** Variants were partitioned into training and validation sets using family-level metadata to prevent data leakage and ensure evolutionary independence between splits.
- **Standardization and Imputation:** Features were centered and scaled using `StandardScaler`, with mean-value imputation applied to any missing data points.
- **PCA Projection:** Principal Component Analysis (PCA) was utilized to reduce dimensionality while retaining 99% of the cumulative explained variance.

### Supervised Classification and Optimization

A deep multi-layer perceptron (MLP) was trained to predict protein stability as a binary classification task (Δ*G >* 3 kcal/mol). We utilized `Optuna` for 300 trials of Bayesian hyperparameter optimization, tuning architecture depth, layer widths, dropout rates, and learning schedules. The final model weights were selected based on the peak validation F1-score:

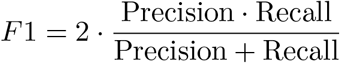

Performance was rigorously evaluated on independent test sets, utilizing confusion matrices and class-specific recall to validate the model’s generalizability across diverse protein families.

## Ablation models

### Topological Null Model: Random Edge Rewiring

To isolate the contribution of protein network topology from simple spatial proximity, we generated randomized graphs using a Topology Noising strategy. This method produces a null model where the protein fold and edge weight distribution are conserved, but the specific biological connectivity is destroyed.

### Conservation of Graph Invariants

The randomization process was constrained to maintain the first-order properties of the native protein graph *G* = (*V, E*), where *V* represents the set of nodes (amino acid residues) and *E* represents the set of edges (physicochemical interactions). For each input graph, we preserved the node set *V* and its associated spatial attributes (PDB Cartesian coordinates). We extracted the set of original edge weights *W*, where each weight *w ∈ W* corresponds to the strength of a specific bond (e.g., hydrogen, peptide, or hydrophobic) as defined by the physicochemical properties of the residues.

### Algorithmic Implementation of Random Rewiring

The original edge set *E* was entirely purged, effectively removing the biological connectivity while leaving the residues fixed in 3D space. A new, stochastic edge set *E*_*rand*_ was then populated using a batch-sampling approach to achieve an identical edge count |*E*_*rand*_| = |*E*| .

The generation of *E*_*rand*_ followed a uniform sampling distribution:

- **Node Selection:** Pairs of node indices (*u, v*) were sampled from a discrete uniform distribution *U* (0, |*V* |−1), where |*V* | is the total number of residues in the protein.
- **Uniqueness Constraints:** To ensure an undirected, simple graph, self-loops (*u* = *v*) were prohibited. Pairs were sorted such that (*u, v*) ≡ (*v, u*) to prevent duplicate edges.
- **Batch Processing:** Sampling was performed in batches of 1.5 × |*E*| to optimize computational efficiency against collisions, continuing until the target edge cardinality was reached.

### Weight Shuffling and Mapping

To preserve the exact chemical “budget” of the protein while removing structural context, the weight vector *W* was permuted using a Fisher-Yates shuffle. Each shuffled weight 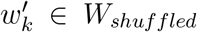 assigned to a corresponding edge *e*_*k*_ ∈ *E*_*rand*_.

The resulting adjacency matrix *A*_*rand*_ for the noised graph is defined as:

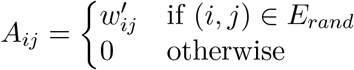

Where 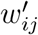 is the randomized weight assigned to the connection between residues *i* and *j*. This ensures that the total graph weight Σ*w*_*ij*_ and the distribution of bond strengths are invariant, but the adjacency relationship between residues is entirely decoupled from their sequence distance or biological proximity. To ensure the reproducibility of this stochastic topology across different job arrays, a global random seed of 42 was applied to the randomization modules.

### Geometric Null Model: Rigid Body Random Walk

To assess the impact of the native 3D fold on network topology, we implemented a **Geometric Random Walk** protocol. This approach generates a non-biological, “random coil” spatial configuration of the protein while preserving the local rigid-body geometry of individual amino acids and their primary sequence order.

### Stochastic Backbone Trajectory

The global trajectory of the protein backbone was modeled as a self-avoiding-like random walk in ℝ3. Let **p**_*i*_ denote the Cartesian coordinate vector (*x, y, z*) of the *α*-carbon (*Cα*) for the *i*-th residue in a sequence of total length *N* . The coordinates were generated iteratively according to the following process:

1. **Initialization:** The first residue in the sequence was anchored at the origin, where **p**_1_ = (0, 0, 0).
2. **Directional Sampling:** For each subsequent residue *i* ∈ > { 2, …, *N* }, a random unit direction vector 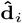 was sampled from a uniform distribution on a sphere. This was achieved via Gaussian normalization:

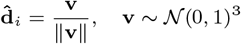

where **v** is a 3D vector of independent standard normal samples.
3. **Step Displacement:** To simulate backbone flexibility, a scalar step size *s*_*i*_ was sampled from a uniform distribution *U* (2.0, 4.0) Å. The position of the current residue was then defined as:

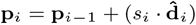

where **p**_*i−*1_ is the position of the preceding residue in the sequence.

### Rigid-Body Side-Chain Transformation

To maintain the chemical integrity of individual residues, each amino acid was treated as a rigid body. Let *{***r**_*i,j*_*}* be the set of relative coordinate vectors for all atoms *j* in residue *i*, defined as the distance from the native *Cα* position in the original PDB structure. For each residue *i*, a unique random rotation matrix **R**_*i*_ *∈ SO*(3) was generated. The final transformed absolute coordinates **x**_*i,j*_ for every atom *j* in the null model were calculated as:

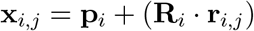

In this equation, **p**_*i*_ represents the new *Cα* position from the random walk, and **R**_*i*_ . **r**_*i,j*_ represents the randomly reoriented side-chain atoms. This ensures that internal bond lengths and angles within each residue remain identical to the biological source, while their orientation relative to the chain and the global fold is randomized.

### Geometry-Dependent Edge Re-calculation

Following the spatial randomization, the network edges *E*_*rw*_ were re-calculated *de novo* based on the new coordinate set **x**_*i,j*_. Unlike the topological noising where edges were sampled stochastically, here the edges are a deterministic function of the new geometry:

- **Sequence Edges:** Peptide bonds were maintained between adjacent residues (*i, i* + 1) in the sequence.
- **Non-Covalent Interactions:** Edge types such as hydrogen bonds, hydrophobic clusters, and disulfide bridges were assigned to the new edge set *E*_*rw*_ only if the transformed atomic coordinates **x**_*i,j*_ met the specific distance thresholds *d* defined in the native interaction model (e.g., *d <* 3.5 Å for hydrogen bonds).

The resulting graph *G*_*rw*_ = (*V, E*_*rw*_), where *V* is the set of residues and *E*_*rw*_ is the new set of geometry-derived edges, represents a chemical polymer with the same composition as the native protein but lacking an evolved tertiary structure. Reproducibility was ensured by applying a global random seed of 42 to the `numpy.random` and `scipy.spatial.transform` modules.

### Geometric Null Model: Bounding Box Randomization (Random box model)

To evaluate the significance of the spatial distribution without the influence of sequence connectivity, we implemented a **Bounding Box Randomization** strategy. This model creates a gas-like distribution of residues that preserves the overall volume and density of the native protein but eliminates all structural and sequential context.

### Preservation of Global Spatial Envelope

The randomization is constrained by the original spatial dimensions of the native protein to ensure the null model occupies a comparable volume. For a protein with *N* residues, the original Cartesian coordinates (*x*_*i*_, *y*_*i*_, *z*_*i*_) were used to define a bounding box:

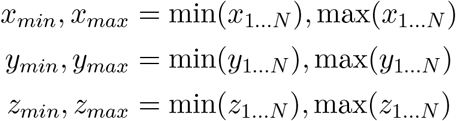

These limits define the domain Ω = [*x*_*min*_, *x*_*max*_] *×* [*y*_*min*_, *y*_*max*_] *×* [*z*_*min*_, *z*_*max*_], ensuring that the randomized residues are contained within the same macroscopic 3D envelope as the biological structure.

### Uniform Spatial Sampling

Each residue *i* ∈ *{*1, …, *N}* is assigned a new position 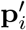 sampled from a continuous uniform distribution across the three spatial axes:

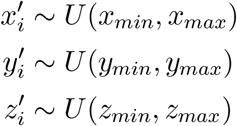

Unlike the random walk model, the distance between residue *i* and *i* + 1 is not constrained, effectively shattering the polypeptide chain into a set of independent points in space. This isolates the effect of residue density and chemical composition from the effects of backbone topology.

### Graph Reconstruction and Feature Extraction

In this null state, the “PDB position” attribute (pos*i*) is updated to the randomized coordinates:

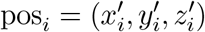

This updated position serves as the basis for all downstream spatial feature calculations.

## SHAP Analysis

To interpret the contributions of individual features to the predictive performance of the deep learning models, we utilized **SHAP (SHapley Additive exPlanations)**. SHAP is a game-theoretic approach that assigns each feature an importance value (a Shapley value) for a particular prediction, representing the change in the expected model output when that feature is observed versus when it is missing.

Given the computational complexity of the deep learning architectures (Cases 1–6), we employed the `KernelExplainer` with a K-means summary of the training data (20 centroids) used as the background distribution. This allowed for an efficient estimation of feature impact across a unified test set (*N ≈*1200 samples across all cases).

### Quantitative Metrics for Feature Importance

For each feature *i*, the primary metric of importance is the **Mean Absolute SHAP value**, defined as:

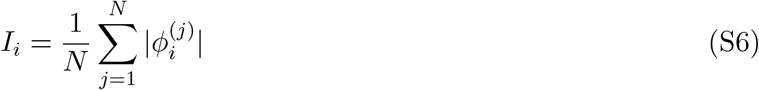

Where 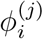 is the SHAP value for feature *i* in sample *j*. Based on this, we derived two secondary metrics to evaluate the contribution of different feature engineering categories:

1. **Importance Share (%)**: The cumulative impact of a specific category *C* relative to the total model signal. This indicates the total “weight” assigned to a specific pipeline stage.

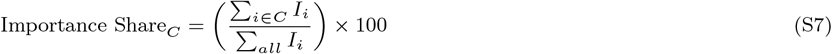
2. **Average Potency**: The mean importance of features within a category, used to identify which stages provide the highest quality signal per individual feature (also referred to as average importance per feature).

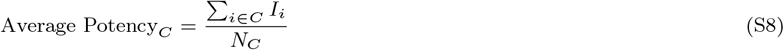

Where *N*_*C*_ is the total number of features contained within category *C*.

### Categorization Framework

To analyze the biological and structural logic of the model, the 294 features were bifurcated through three distinct categorization lenses:

- **Pipeline Stage**: Features were grouped by their mathematical origin into *Connectivity-based* (topology), *Geometry-based* (spatial metrics), and *Projection-based* (centroid/volume metrics).
- **Multi-Resolution Subnetworks**: Features were grouped by their subnetwork index ( 0 to 5). This identifies which structural resolution threshold contains the most critical predictive information.
- **Intra-vs. Inter-Subnetwork**: Features were classified as *Intra-subnetwork* (representing local topological slices) or *Inter-subnetwork* (representing global chemical interactions such as Hydrophobic-Polar pairings).

---

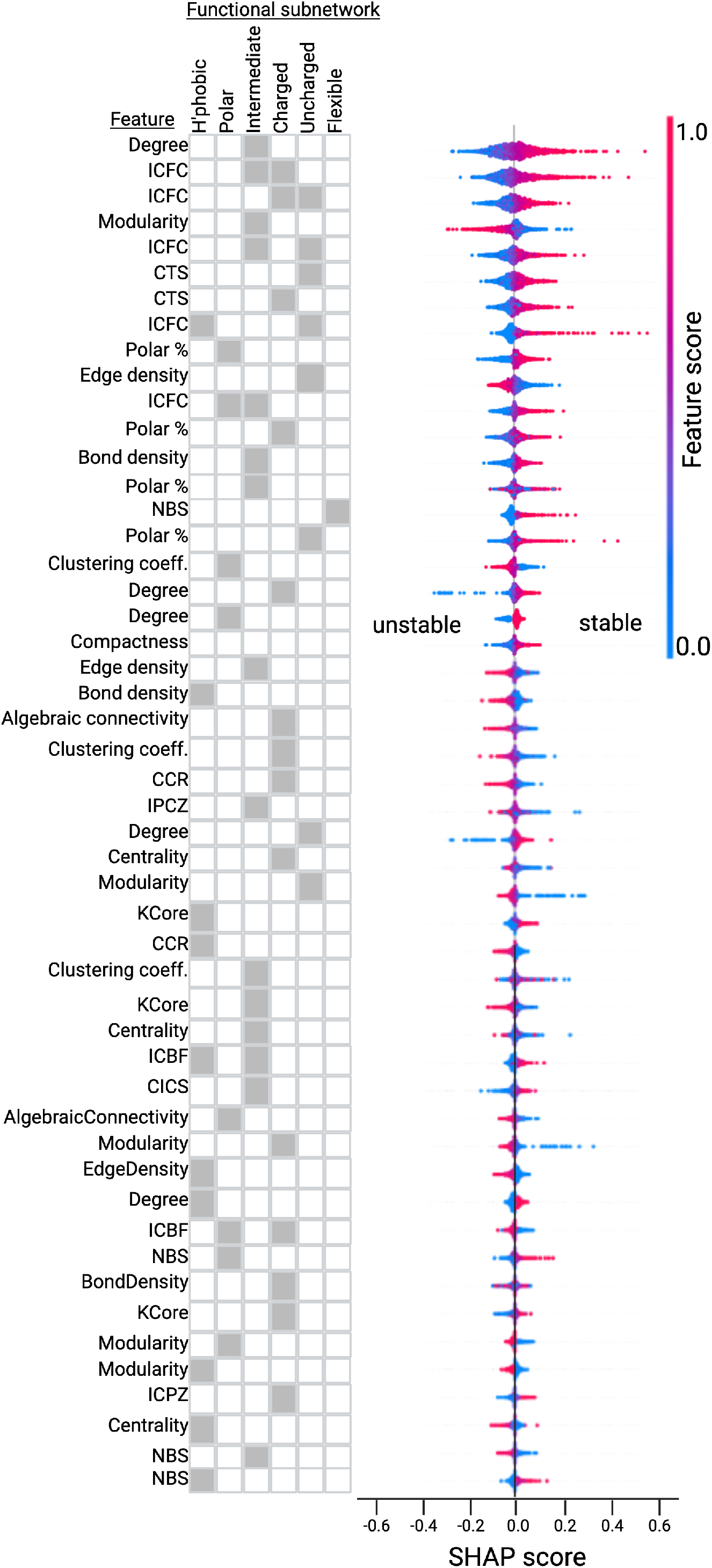

## References

1. Christian B Anfinsen. Principles that govern the folding of protein chains. Science, 181(4096):223–230, 1973.

2. Christopher M Dobson. Protein folding and misfolding. Nature, 426(6968):884–890, 2003.

3. John Jumper, Richard Evans, Alexander Pritzel, et al. Highly accurate protein structure prediction with AlphaFold. Nature, 596(7873):583–589, 2021.

4. Po-Ssu Huang, Scott E Boyken, and David Baker. The coming of age of de novo protein design. Nature, 537(7620):320–327, 2016.

5. Debora S Marks, Lucy J Colwell, Robert Sheridan, et al. Protein 3d structure computed from evolutionary sequence variation. PloS One, 6(12):e28766, 2011.

6. Ken A Dill and Justin L MacCallum. The protein-folding problem, 50 years on. Science, 338(6110):1042–1046, 2012.

7. Alan Fersht. Structure and mechanism in protein science: a guide to enzyme catalysis and protein folding. Macmillan, 1999.

8. Nobuhiko Tokuriki and Dan S Tawfik. Stability effects of mutations and protein evolvability. Current Opinion in Structural Biology, 19(5):596–604, 2009.

9. Ken A Dill. Dominant forces in protein folding. Biochemistry, 29(31):7133–7155, 1990.

10. Yaakov Levy and Jose N Onuchic. Water and proteins: A love–hate relationship. Proceedings of the National Academy of Sciences, 103(21):8117–8122, 2006.

11. Frances H Arnold. Directed evolution: Bringing new chemistry to life. Angewandte Chemie International Edition, 57(16):4143–4148, 2018.

12. Cyrus Levinthal. How to fold graciously. In Mössbauer spectroscopy in biological systems, pages 22–24, 1969.

13. Rebecca F Alford, Andrew Leaver-Fay, Javi R Jeliazkov, et al. The Rosetta all-atom energy function for macromolecular modeling and design. Journal of Chemical Theory and Computation, 13(6):3031–3048, 2017.

14. Joshua Meier, Roshan Rao, Robert Verkuil, et al. Language models enable zero-shot prediction of the effects of mutations on protein function. Proceedings of the National Academy of Sciences, 118(44):e2109441118, 2021.

15. John Ingraham, Vikas Garg, Regina Barzilay, and Jaakkola Tommi. Generative models for graph-based protein design. In Advances in Neural Information Processing Systems, volume 32, 2019.

16. Markus Reichstein, Gustau Camps-Valls, Bjorn Stevens, et al. Deep learning and process understanding for data-driven earth system science. Nature, 566(7743):195–204, 2019.

17. Palash Sethi and Juannan Zhou. An interpretable neural network unveils higher-order epistasis in large protein sequence-function relationships. bioRxiv, 2024.

18. Enrique Vargas-Vázquez et al. De novo design and stability of miniaturized protein scaffolds. Biomolecules, 14(7):775, 2024.

19. Alexander Rives, Joshua Meier, Tomer Sataluri, et al. Biological structure and function emerged from scaling unsupervised learning to 250 million protein sequences. Proceedings of the National Academy of Sciences, 118(15):e2016239118, 2021.

20. Ethan C Alley, Grigory Khimulya, Surojit Biswas, et al. Unified rational protein engineering with sequence-based deep representation learning. Nature Methods, 16(12):1315–1322, 2019.

21. Zeming Lin, Halil Akin, Roshan Rao, et al. Evolutionary-scale prediction of atomic-level protein structure with a language model. Science, 379(6637):1123–1130, 2023.

22. Justas Dauparas, Ivan Anishchenko, Nathaniel Bennett, et al. Robust deep learning–based protein sequence design using ProteinMPNN. Science, 378(6618):442–451, 2022.

23. Surojit Biswas, Grigory Khimulya, Ethan C Alley, et al. Toward machine-guided design of proteins. Nature Methods, 18(5):489–496, 2021.

24. Patrick Bryant, Gabriele Pozzati, and Arne Elofsson. Improved prediction of protein-protein interactions using AlphaFold2. Nature Communications, 12(1):6912, 2021.

25. Henry Dieckhaus et al. ThermoMPNN: structural foundation model for predicting stability changes upon mutation. bioRxiv, 2024.

26. Kotaro Tsuboyama, Justas Dauparas, Jonathan Chen, Elodie Laine, Yasser Mohseni Behbahani, Jonathan J Weinstein, Niall M Mangan, Sergey Ovchinnikov, and Gabriel J Rocklin. Mega-scale experimental analysis of protein folding stability in biology and design. Nature, 620(7973):434–444, 2023.

27. Chloe Hsu, Hunter Nisonoff, Colleen Fannjuan, et al. Learning protein fitness models from evolutionary and assay-labeled data. Nature Biotechnology, 40(7):1114–1122, 2022.

28. Kevin Gao et al. Limitations of sequence-only protein language models in the design of de novo protein-protein interactions. bioRxiv, 2023.

29. Cynthia Rudin. Stop explaining black box machine learning models for high stakes decisions and use interpretable models instead. Nature Machine Intelligence, 1(5):206–215, 2019.

30. Julius Adebayo, Justin Gilmer, Michael Muelly, et al. Sanity checks for saliency maps. In Advances in neural information processing systems, volume 31, 2018.

31. Amirata Ghorbani, Abubakar Abid, and James Zou. Interpretation of neural networks is fragile. In Proceedings of the AAAI Conference on Artificial Intelligence, volume 33, pages 3681–3688, 2019.

32. Albert-László Barabási and Zoltán N. Oltvai. Network biology: understanding the cell’s functional organization. Nature Reviews Genetics, 5(2):101–113, 2004.

33. Debnath Sengupta, Sankar Kundu, and Saraswathi Vishveshwara. Network analysis of protein structures. Proceedings of the National Academy of Sciences (PNAS), 101(34):12526–12531, 2004.

34. Ernesto Estrada. Topological structural analysis of proteins. can individual residues tell us anything about protein folding? Bioinformatics, 18(2):301–307, 2002.

35. John Jumper et al. Highly accurate protein structure prediction with alphafold. Nature, 596(7873):583–589, 2021.

36. Zeming Lin et al. Evolutionary-scale prediction of atomic-level protein structure with a language model. Science, 379(6637):1123–1130, 2023.

37. G. Amitai, A. Shemesh, E. Sitbon, M. Shklar, D. Netanely Venger, and S. Pietrokovski. Network analysis of protein structures identifies functional residues. Journal of Molecular Biology, 344(4):1135–1146, 2004.

38. Damiano Piovesan, Giovanni Minervini, and Silvio C. E. Tosatto. The ring 2.0 web server for high quality residue interaction networks. Nucleic Acids Research, 44(W1):W367–W374, 2016.

39. Guillaume Brysbaert and Marc F. Lensink. Centrality measures in residue interaction networks to highlight amino acids in protein–protein binding. Frontiers in Bioinformatics, 1, 2021.

40. J. Dauparas et al. Robust deep learning–based protein sequence design using proteinmpnn. Science, 378(6615):49–56, 2022.

41. Mohammed AlQuraishi. Machine learning in protein structure prediction. Current opinion in chemical biology, 65:1–8, 2021.

42. Ruidong Wu, Fan Ding, Rui Wang, Rui Shen, Xiwen Zhang, Shaohuai Luo, Chi-Hua Su, Zhewen Wu, Qi Xie, Biqing Pan, et al. High-resolution de novo structure prediction from protein sequences. bioRxiv, pages 2022–07, 2022.

43. Vincent D. Blondel, Jean-Loup Guillaume, Renaud Lambiotte, and Etienne Lefebvre. Fast unfolding of communities in large networks. Journal of Statistical Mechanics: Theory and Experiment, 2008(10):P10008, 2008.

44. Thomas M. J. Fruchterman and Edward M. Reingold. Graph drawing by force-directed placement. Software: Practice and Experience, 21(11):1129–1164, 1991.

45. Josh Abramson, Jonas Adler, Jack Dunger, Richard Evans, Tim Green, Alexander Pritzel, Olaf Ronneberger, Lindsay Willmore, Andrew J. Ballard, Joshua Bambrick, et al. Accurate structure prediction of biomolecular interactions with AlphaFold 3. Nature, 630(8016):493–500, 2024.

46. Jack Kyte and Russell F. Doolittle. A simple method for displaying the hydropathic character of a protein. Journal of Molecular Biology, 157(1):105–132, 1982.

47. Richard Grantham. Amino acid difference formula to help explain protein evolution. Science, 185(4154):862–864, 1974.

48. Ludovica Montanucci, Emidio Capriotti, Yotam Frank, Nir Ben-Tal, and Piero Fariselli. Ddgun: an untrained method for the prediction of protein stability changes upon single and multiple point variations. BMC bioinformatics, 20(Suppl 14):335, 2019.

49. Iván Martín Hernández, Yves Dehouck, Ugo Bastolla, José Ramón López-Blanco, and Pablo Chacón. Predicting protein stability changes upon mutation using a simple orientational potential. Bioinformatics, 39(1):btad011, 2023.

50. Ziang Li and Yunan Luo. Generalizable and scalable protein stability prediction with rewired protein generative models. Nature Communications, 2025.

51. Corrado Pancotti, Silvia Benevenuta, Giovanni Birolo, Virginia Alberini, Valeria Repetto, Tiziana Sanavia, Emidio Capriotti, and Piero Fariselli. Predicting protein stability changes upon single-point mutation: a thorough comparison of the available tools on a new dataset. Briefings in Bioinformatics, 23(2):bbab555, 2022.

52. Scott M Lundberg and Su-In Lee. A unified approach to interpreting model predictions. In I. Guyon, U. V. Luxburg, S. Bengio, H. Wallach, R. Fergus, S. Vishwanathan, and R. Garnett, editors, Advances in Neural Information Processing Systems 30, pages 4765–4774. Curran Associates, Inc., 2017.

